# Homoeo-alleles of wheat *GNI2* fuel grain yields across input environments

**DOI:** 10.64898/2026.04.28.721284

**Authors:** Shun Sakuma, Matteo Bozzoli, Guy Golan, Manar Makhoul, Cristian Forestan, Kenan Tan, Ahmed Raza Khan, Francesco De Sario, Sara Giulia Milner, Giuseppe Sciara, Chuny Liu, Elisabetta Frascaroli, Fumitaka Abe, Goetz Hensel, Jia-Wu Feng, Martin Mascher, Karim Ammar, Susanne Dreisigacker, Mikiko Kojima, Masanori Okamoto, Roberto Tuberosa, Silvio Salvi, Rod Snowdon, Marco Maccaferri, Thorsten Schnurbusch

## Abstract

Grain yield in wheat is frequently driven by pre-anthesis growth and how carbon is directed to reproductive parts, determining the potential size of the grain-producing sink. However, the genetic mechanisms controlling this carbon allocation remain unclear. In this study we discovered a series of loci in tetraploid and hexaploid wheats that, when combined, confer an average grain yield advantage in hexaploid winter wheat of approx. 5-7% under high-input farming while approx. 10-15% under low-input conditions. The responsible gene is *GRAIN NUMBER INCREASE 2* (*GNI2*), ancestral to its duplicate *GNI1*, and represented by the homoeo-allelic series *GNI-A2*, *GNI-B2*, and *GNI-D2*. *GNI1* and all *GNI2* copies additively affect floral growth and fertility by modulating reproductive allocation and harvest index. Herewith, we describe a series of grain yield-relevant *GNI2* homoeo-alleles with a proven track record for being beneficial in both high- and low-input environments. Their deployment offers a sustainable pathway to raise global grain yields in the future.

## Introduction

The stability and accessibility of primary production and food supply are essential for the prosperity and functioning of our global civilization, establishing food security as a vital cornerstone of the modern era^1^. The global demand for sources of our staple foods, such as rice (*Oryza sativa* L.), wheat (*Triticum* sp.), and maize (*Zea mays* L.), is projected to increase by approx. 50% in 2050^2,3^. Achieving this sustainably despite global climate change, soil loss, and geo-political instabilities remains one of the pivotal factors^1^. Considering that the global grain yield average of wheat is approx. 3.5 t/ha^4^, sustainable increases in developed, low and middle income countries are thus of foremost significance^3^. Empirically, plant breeding has a longstanding history of genetically enhancing crop yields, thereby providing evidence that wheat grain yields are primarily raised by reproductive allocation, enhancing grain number per unit area^5–8^; and here, mainly through more grains per spike^9,10^. However, increasing grain number is a many-faceted process^11^ and not entirely genetically driven since it is similarly affected by the crops’ phenology, soil, and weather conditions, making it a complex, quantitative, and multi-factorial trait. Therefore, annual average global grain yield gains in wheat are low and range from 0.8 to 1.5% per year, while they are usually lower or even stagnating in high-yielding regions such as Western Europe^12^.

The grain-bearing organs in wheat are called florets (grass flower); they are produced in small inflorescences, termed spikelets, which represent the basic reproductive units of all grasses^13^. Wheat plants produce spikelets that are alternate-distichously attached to the rachis, i.e. the axis of their spike-type inflorescences. Thus, rachis and rachilla, i.e. the spikelet axis, are structurally supporting tissues for the florets through which most of the assimilate supply is conducted. To determine natural variation for floret and grain number per spikelet in wheat, late phenotyping of spikelets is required, i.e. during grain-fill or after physiological maturity. High-fertility genotypes are characterized by an extended rachilla activity resulting in more florets per spikelets, eventually producing more grain. Floret and grain number per spikelet appear as quantitative traits that are affected by multiple genetic loci^14^. Since grain numbers in wheat are mainly driven during pre-anthesis growth, carbon allocation to reproductive structures during this critical period promotes floral growth and increases potential sink size^11,15^. In many cereals, however, apical floral primordia are often destined to undergo degeneration during later stages of floral development^16^. Importantly, wheat and barley (*Hordeum vulgare* L.) plants actively reduce their sink size by up to ∼50%^14,17^, most likely as a direct consequence of intra-organ, intra-plant, or inter-plant competitive events for resources in combination with a floral degeneration program, which is adaptive to the growth environment^18–20^. For example, barley spikes lose approx. 15-50% of their most apical floral primordia^21^, representing a significant loss of their grain yield potential. This phenomenon, termed pre-anthesis tip degeneration (PTD), appears to be a multi-layered process that is largely developmentally programmed^22,23^, while its extent, i.e. how many floral structures will degenerate, is predominantly dependent on genotype and its interactions with the environment^24–26^. Recent molecular investigations of PTD showed that barley *GRASSY TILLERS 1* (*HvGT1*), the ortholog of maize *GT1*^27^, is a prominent modulator of spike PTD by inhibiting tip growth and floral differentiation^23^. *HvGT1* belongs to the class I homeodomain leucine zipper protein family of transcriptional regulators that have a well-known function as growth repressors^28^. Notably, wheat *GRAIN NUMBER INCREASE 1* (*GNI1*), a close paralog of *HvGT1*, similarly exerts an inhibitive growth function on the most apical floral primordia, suggesting that barley and wheat might share a common genetic and molecular basis for PTD. Most importantly, the reduced function allele of *gni-A1105Y* leads to higher floral fertility and grain set in wheat plants and increased plot grain yields^9^.

In this study we discovered a series of homoeo-loci in tetraploid and hexaploid wheats that, when combined with *gni-A1105Y*, confer an average grain yield advantage in hexaploid winter wheat of approx. 5-7% under high-input conditions while approx. 10-15% under low-input farming. The responsible gene *GRAIN NUMBER INCREASE 2* (*GNI2*), represented by the homoeo-allelic series *GNI-A2*, *GNI-B2*, and *GNI-D2*, is ancestral to its duplicate *GNI1*. Both additively affect floral growth and fertility through modulating source-sink relations, reproductive allocation and harvest index. Herewith, we identify a series of grain yield-relevant *GNI2* homoeo-alleles in wheat that have a proven track record for being beneficial in both high- and low-input environments, offering a sustainable pathway to further raise global grain yields in future.

## Results

### Discovery of *GNI2* homoeo-allelic loci in tetraploid wheat with increased floret and grain number per spikelet

To obtain genetic access to high-fertility genotypes that confer extended rachilla activity and more florets per spikelet, we used contrasting wild and domesticated tetraploid wheat accessions in our investigations. Selected parental lines of our populations showed heritable genetic variation for the number of fertile florets and grains per spikelet or spike (Figure 1A-C). We then analyzed multi-year field and greenhouse experiments for spikelet fertility and grain number per spike and found two consistent quantitative trait loci (QTL) in homoeo-allelic regions on the short arms of chromosomes 2A and 2B (Figure 1D-E; Suppl Table S1). To identify the favorable allele underlying *QGns.ubo-2A*, we used recombinants from a total of 1,550 individual F5 plants derived from a cross between *cvs*. Relief and Iride (Figure 1C) segregating for the floral fertility trait. Fine-mapping *QGns.ubo-2A* located the interval within a 3.9-Mb region, harboring 32 high-confidence genes in the Svevo reference genome (Suppl Table S2). The interval includes *TRITD2Av1G066050* encoding for a homeodomain leucine zipper class I (HD-Zip I) transcription factor, which is a paralog of wheat *GNI1*^9^ and orthologous to barley *Hox2*^29,30^. A comparison of the two parental alleles revealed a 3,219 bp-deletion that includes the entire open-reading frame of *TRITD2Av1G066050* (hereafter *GNI-A2*) in Rascon/2*Tarro, *cv*. AltarC84, and Iride (Figure 1D) resulting in the *gni-A2del* deletion allele. For the syntenic 2BS region we used a cross between wild (Zavitan) and domesticated wheat (*cv.* Svevo). Due to the putative homoeo-allelic relationship between the two detected QTL, we followed a comparative approach and found that the true homoeolog of *GNI-A2*, i.e. *TRITD2Bv1G077460* (hereafter *GNI-B2*), was directly located under the QTL peak. Sequence analysis of *TRITD2Bv1G077460* revealed a frameshift mutation at nucleotide position 298 prior to the homeodomain (Figure 1E) leading to a *loss-of-function* allele in Svevo (*gni-B2298_del*). We then validated and confirmed the higher floret number phenotype under greenhouse conditions in a mutant of Svevo, which already naturally carries the *gni-B2298_del* allele (Figure 1E) and the reduced-function *gni-A1105Y* allele, but also has a newly induced mutation in *GNI-A2* leading to an amino acid substitution (L116F; Figure 1D; Suppl Figure S1) shortly after the homeodomain. Although showing more florets per spikelet under greenhouse conditions, this triple mutant T201 (*gni-A1105Y*/*gni-A2116F*/*gni-B2298_del*) in tetraploid wheat failed to have more grains per spike (Suppl Figure S1), most likely due to high mutational background load. For hexaploid wheat, we pursued a CRISPR-based knock-out (KO) approach of all three homoeologous *GNI2* copies (hereafter referred to as *GNI2s*) using the US-derived *cv*. Fielder. From T2-derived plants, we generated all seven possible single-, double-, and triple-KO lines (Figure 2) and screened them phenotypically under controlled greenhouse conditions against T2-derived plants carrying only wild-type alleles for *GNI2s* (AABBDD). Our analyses revealed that all pot-grown greenhouse plants appeared phenotypically very similar, while only plants carrying the triple-KO mutations (aabbdd) had more florets per spikelet and an extended rachilla phenotype (Figure 2C); however, wild-type, single, and double mutants displayed wild-type-like floral features with reduced floral numbers. Our greenhouse data thus suggests that high-fertility genotypes in hexaploid wheat can reliably be identified based on higher floret numbers per spikelet and the extended post-anthesis rachilla phenotype.

**Figure 1.**
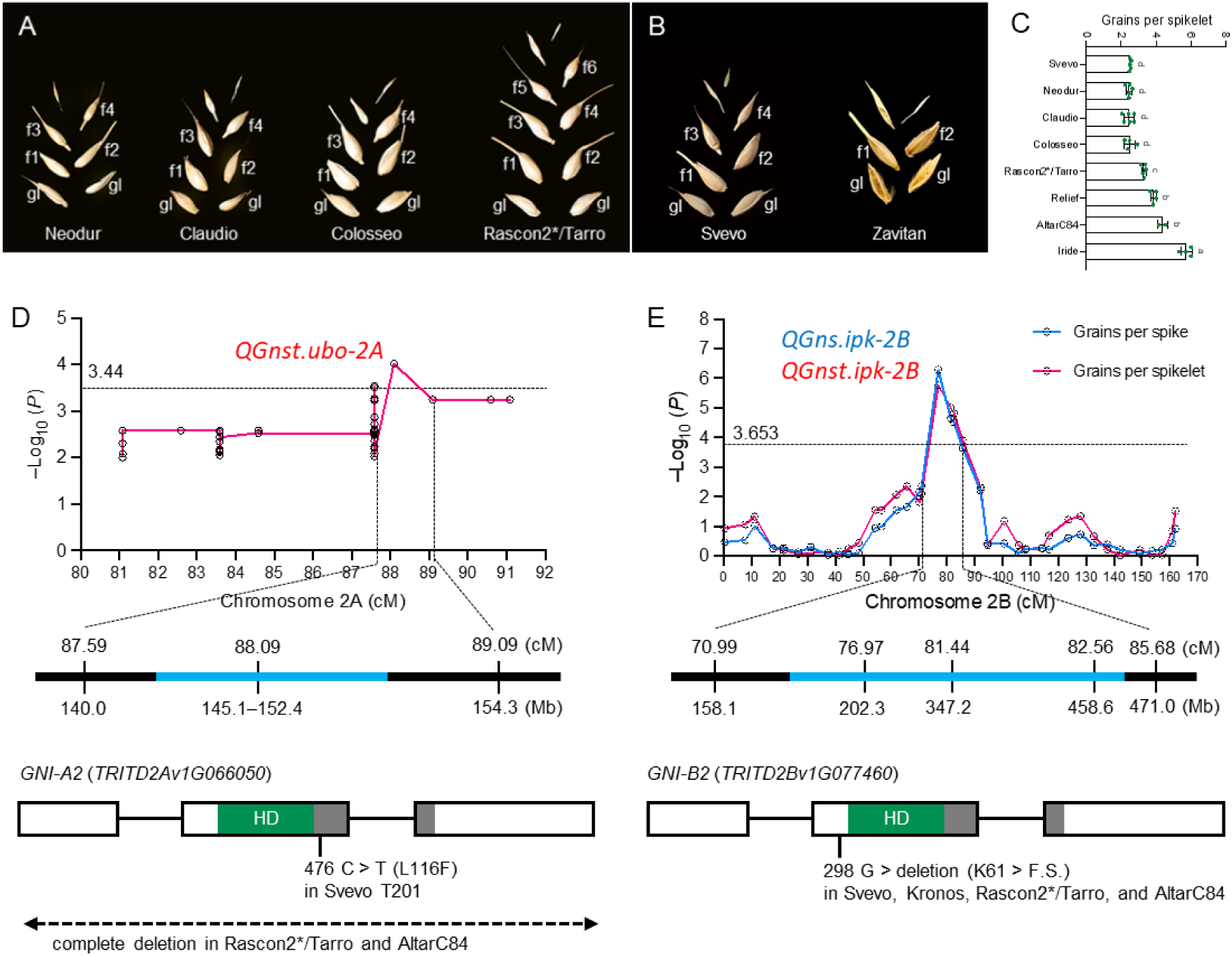
Identification and positional cloning of two *GNI2* homoeo-loci for improved floral enhancement. (A) Spikelet structure of the parental lines in the NCCR mapping population; gl, glume; f, floret. (B) Spikelet structure of Svevo and Zavitan. (C) Grain number per spikelet. Letters are used to indicate where mean values differed from one another significantly (P< 0.05) as determined by a one-way ANOVA with Tukey’s multiple comparison test. Scale bar = 5 cm. (D) QTL mapping of *GNI-A2* in NCCR population. <E) QTL mapping of *GNI-B2* in Zavitan x Svevo population. The horizontal dashed lines represent a threshold LOD value. Sky blue bars indicate the candidate genetic intervals. Boxes indicate exons. The grey boxes indicate a leucine zipper motif. HD, homeodomain. F.S., frameshift.

**Fig 2.**
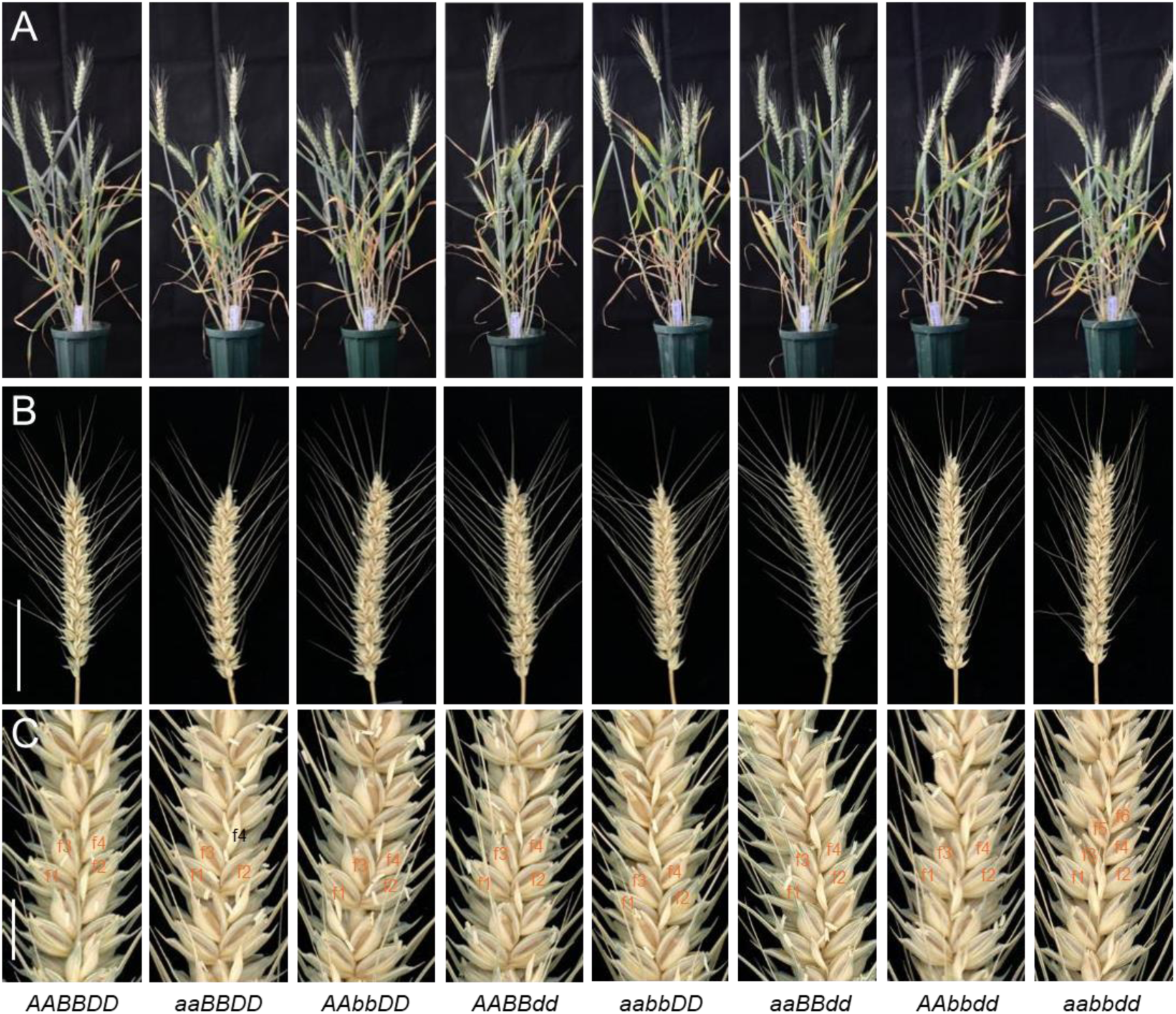
*GNI2* single-, double-, and triple- homoeo-allelic loss-of-function mutants in hexaploid wheat under controlled greenhouse conditions. (A) Plant stature. (B) Spike structure. (C) Floret number per spike. Scale bars = 5 cm in (B) and 1 cm in (C).

### Natural variation at *GNI2* homoeo-loci is associated with spike fertility and grain yield

From our previous work on *GNI1*, we knew that the favorable high-yielding *gni-A1105Y* allele had been selected for during crop evolution in tetraploid durum wheats (*T. durum* L.) and nowadays reached an allele frequency of ≥95%^9^. We therefore sought to estimate the allele frequencies and haplotype diversity of the two newly identified homoeo-loci, *GNI-A2* and *GNI-B2*, in 96 diverse tetraploid wheat accessions^31^, including wild and domesticated forms as well as diverse subspecies (Suppl Figure S2A-E). For *GNI-A2*, we detected 11 mutations in the 5’/3’ UTRs and CDS corresponding to 10 SNPs and 1 deletion (Supp. Info Figure 1). Here, Hap-A1 corresponds to the wild-type genotypes similar to Svevo, and it is well represented in all the 96 selected accessions (Suppl Figure S2A). The *gni-A2del* deletion allele (Hap-A6) identified in AltarC84 had a highly significant positive effect for grains per spikelet in the same panel (Suppl Figure S2C) but at a very low frequency of ∼5% (4 out of 96) and only occurred in known high-fertility cultivars, such as CIMMYT’s AltarC84, and its derivatives Iride (ITA), DBA-Aurora (AUS) and Saragolla (ITA). Importantly, all four genotypes consistently exhibit the high-fertility floral phenotype and share the allele combination *gni-A1105Y*, *gni-A2del* and *gni-B2298_del*. Moreover, since they are all direct descendants of AltarC84, this is line anecdotal evidence for high fertility in AltarC84. The AltarC84 Hap-A6 was then tracked in a more recent set of 456 advanced durum lines representative of the CIMMYT breeding program and was detected in 86 lines, or with a frequency of 19%, consistent with the favorable allele effect on grain yield (Suppl Data S7-8; Supp. Info Figure 2).

The loss-of-function *gni-B2298_del* allele was only found in domesticated forms and had approx. 45% allele frequency in durum wheat (Suppl Figure S2E; Suppl Table S3). However, haplotype analysis of the Hap-B1 in a diverse panel of 96 tetraploid wheat accessions^31^ did not provide a significant effect for more grains per spikelet (Suppl Figure S2B, D), suggesting that the combination of the two alleles, *gni-A1105Y* and *gni-B2298_del*, in most of these accessions is not sufficient for exhibiting the high-fertility post-anthesis floral phenotype.

To assess nucleotide diversity in hexaploid elite germplasm, we utilized two bread wheat (*T. aestivum* L.) panels that provide historical wheat diversity over the last ∼40-50 years of European breeding progress—i.e. BRIWECS^10^ and GABI WHEAT^32^. From both data sets we extracted plot grain yield data over multi-year, multi-site combinations, including yield-component traits such as grain number per spike, harvest index etc., for our haplotype association analyses. Our resequencing and haplotyping efforts among the two panels (210 indiv. GABI WHEAT and 191 BRIWECS) revealed two alleles for *GNI-A2*, three haplotypes for *GNI-B2*, and two alleles for *GNI-D2* (Figure 3A-B; Suppl Figure S3; Suppl Table S4-6). For *GNI-A2*, cultivars carrying the variant 85G (frequency ∼18%) grown under field conditions in multi-year, multi-site combinations produced more grains per spike (*P*=0.004 in BRIWECS population) and higher plot grain yield (*P*=0.04 in GABI; Fig 3E and 3G). For *GNI-B2*, cultivars carrying the variant 23V/71G (∼27%) had increased grains per spike (*P*=0.007) and enhanced plot grain yield (Fig 3F, 3H and J) in the BRIWECS (*P*=0.005) and the GABI WHEAT panel (*P*=0.03). *Hap3*, an apparent recombination between Hap1 and Hap2, carrying 23A and 71G variants, was detected only in two cultivars. *GNI-D2* was rather fixed for one single allele, except for two cultivars with an R42S (Arginine to Serine) amino acid substitution (Suppl Figure S3). In summary, nucleotide variation among the three *GNI2* homoeo-loci in hexaploid wheat followed a well-known pattern thereby showing a significant lack of natural variation within the *DD* genome copy. Greater diversity was found for the *GNI-A2* and *GNI-B2* homoeo-loci; the latter two carried two variants, 85G and 23V/71G, respectively, that showed highly significant genetic associations with enhanced plot grain yield or yield-component traits thereby providing distinct benefits for improved spike fertility and grain yields in bread wheats.

**Figure 3.**
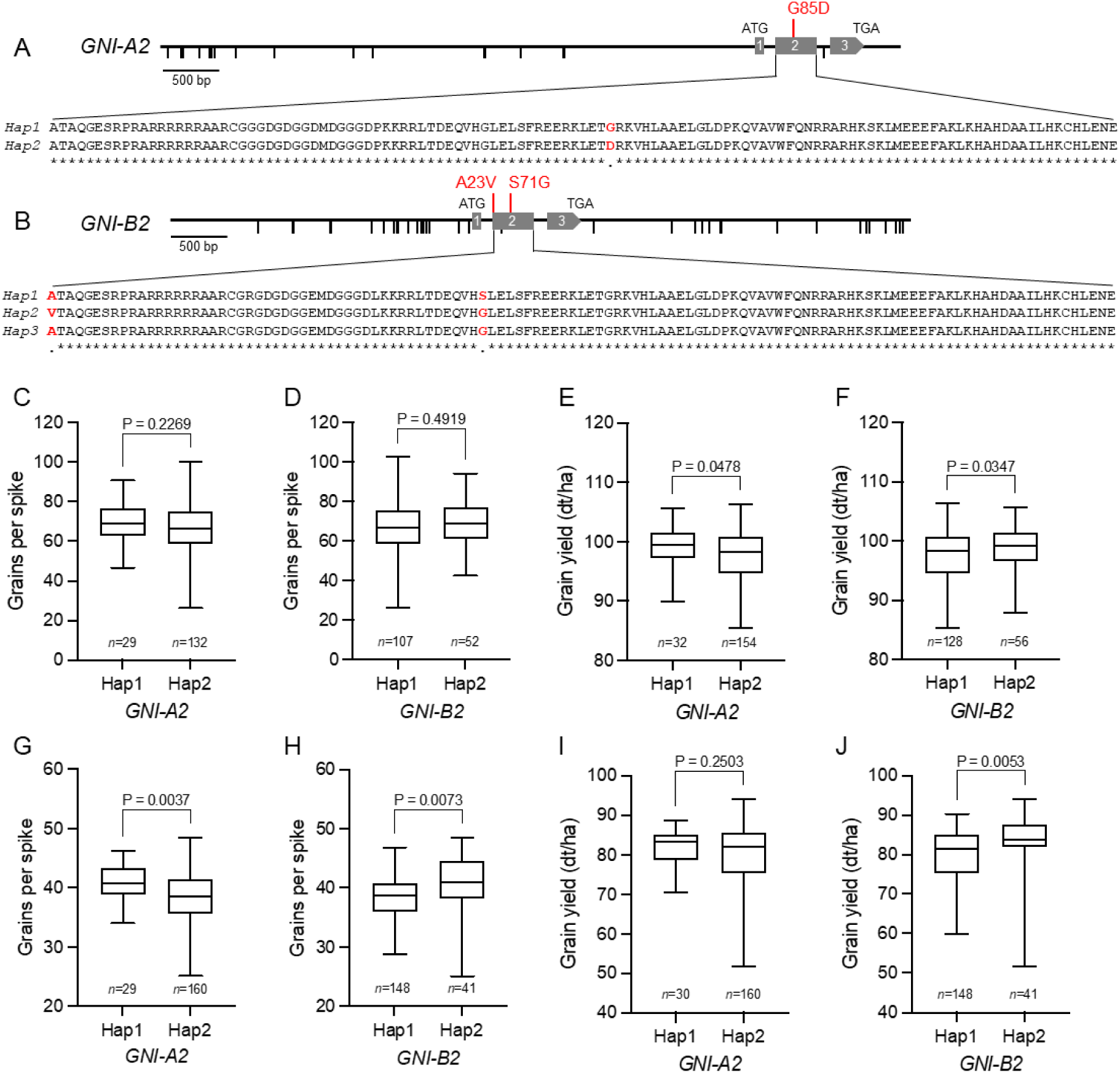
Allelic variation at *GNI2* in bread wheat. (A) *GNI-A2* haplotypes. (B) *GNI-B2* haplotypes. (C-F) A panel of 210 European winter bread wheat cultivars. (G-J) A panel of 199 bread wheat cultivars. The phenotypic data was obtained from work by Guo et al. 2017 (grains per spike), 2018 (grain yield), and Voss-Fels et al. 2019 (grains per spike and grain yield), respectively. A two-tailed Student s *t-* testwas used to determine the significance level. Hap1 refers to the sequences of Chinese Spring.

### *GNI2* homoeo-loci affect plot grain yields in high- and low-input environments

Grain yield is a multifactorial trait that integrates growth conditions with reproductive allocations over the entire life cycle of a wheat crop^7,8,11^. Therefore, to validate our detected genetic associations for individual *GNI2* homoeo-loci we had to establish a solid base for estimating relative grain yield effects on the plot level. To attain these relative grain yield effects for different allele combinations in hexaploid bread wheat cultivars, we took advantage of the available data sets from previous wheat panels—BRIWECS and GABI WHEAT^10,32^. To complement our analysis for *GNI-A2* and *GNI-B2*, we also considered allelic diversity at the *GNI-A1* locus, for which only the known allele *gni-A1105Y* (allele frequency ∼35%) showed positive associations with plot grain yields^9^. For our final analysis, we therefore used allele information from three genetic loci (*GNI-A1*, *GNI-A2*, *GNI-B2*) in combination with the average grain yield data to rank all possible haplotype combinations based upon their grain yield performance. We performed this for the GABI WHEAT panel on the average grain yields (BLUEs) derived from 176 field-grown cultivars from eight environments^32^. For the BRIWECS panel, we used the average grain yields from 186 entries grown in high-input (HiN/F; high nitrogen, fungicide), medium-input (HiN/NoF; high nitrogen, no fungicide), and low-input (LoN/NoF; low nitrogen, no fungicide) conditions from 18, 17, and nine different environments, respectively^10^. Our analysis revealed that across all comparisons and treatments, the three favorable variants at the three loci, i.e. *gni-A1105Y*, *GNI-A285G*, and *GNI-B223V/71G*, were overproportionately enriched in high-yielding cultivars from both panels, which resulted in an average grain yield advantage of approx. 5% compared to the absence of these haplotypes (Figure 4A-D). In all comparisons and treatments, the two top-performing haplotype combinations for grain yield (Hap_1 and Hap_2) contained at least two or three favorable variants with an average grain yield advantage of approx. 8% (Figure 4E). This highlights the critical role of *gni-A1105Y* and *GNI-A285G* for improving grain yields. Under high-input conditions (HiN/F), these two top-performing combinations showed a grain yield advantage ranging from 3-7% for both panels (Figure 4A-B). More importantly, under low-input (LoN/NoF) farming, the average grain yield advantage increased by as much as 10-15% (Figure 4D), indicating that these alleles reliably confer higher grain yields. The superior performance of these haplotype combinations in low-input environments clearly indicates that introducing them into locally adapted germplasm could lead to yield improvements in similar environments worldwide.

**Figure 4.**
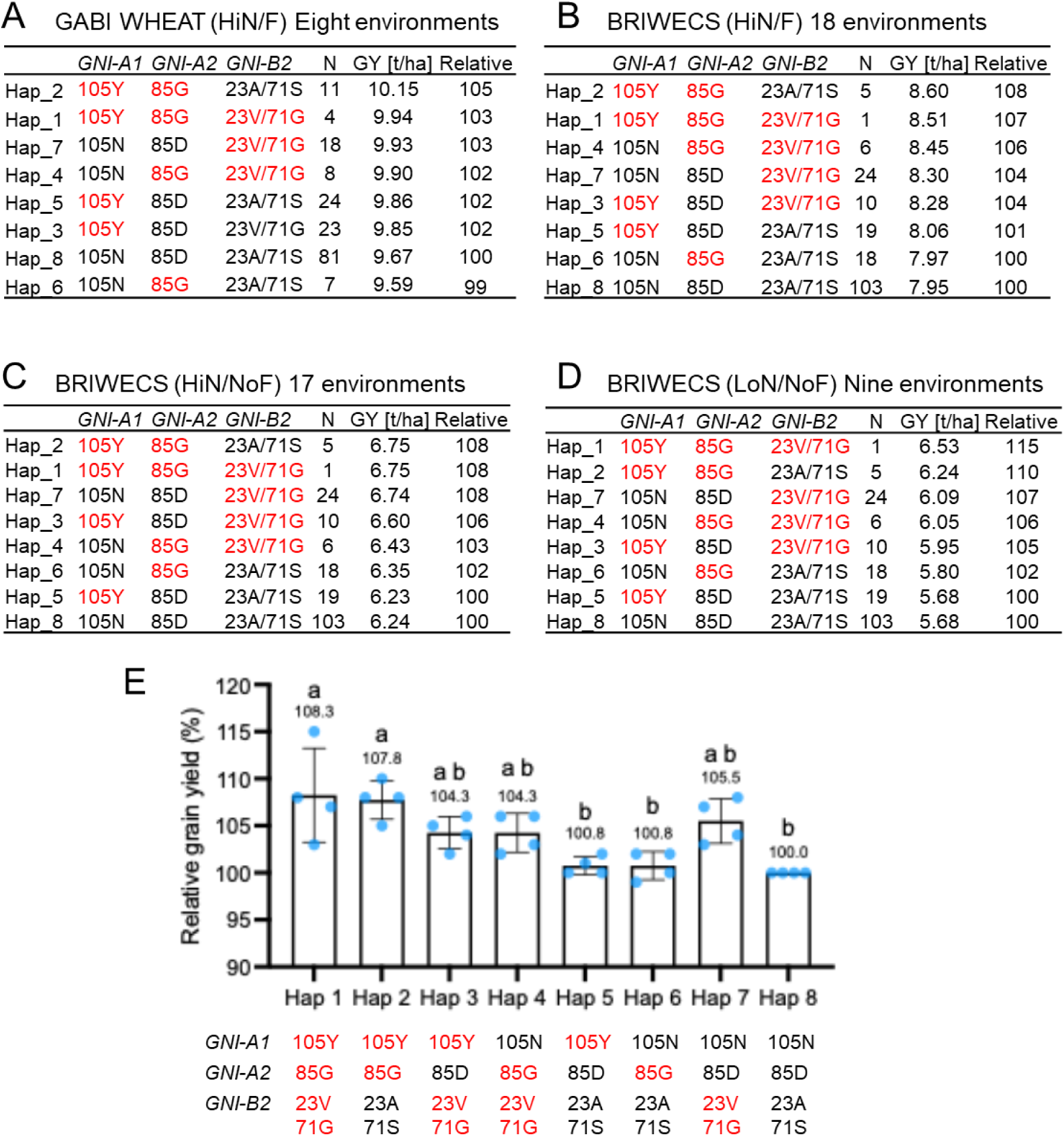
*GNI1* and *GNI2* alleles and haplotypes that contribute to grain yield improvements in hexaploid bread wheat cultivars. Provided are average grain yields (BLUEs, t/ha), average haplotype combination effects from multi-year, multi-site combinations, and the number (N) of cultivars carrying specific combinations. Red color indicates beneficial allele or haplotype. (A) GABI WHEAT panel grown in eight high-input (HiN/F) environments. (B) BRIWECS panel grown in 18 high-input (HiN/F) environments; (C) 17 medium-input (HiN/noF) environments; and (D) nine low-input (LoN/noF) environments. [HiN/F: high nitrogen (N) 220kg/ha, fungicide applications; HiN/noF: high N, no fungicide application; LoN/noF: low N 110kg/ha, no fungicide], (E) Relative haplotype effects across all input environments.

### *gni2* triple mutants raise grain yields in small-plots by modulating reproductive allocation and harvest index

In our previous analyses on natural nucleotide diversity of *GNI2s*, we showed a severe deficiency for *DD* genome variation (Suppl Figure S3); however, this shortcoming also implicated a yet untapped phenotypic potential for enhanced floral fertility for a *DD* genome allele. To explore this in more detail, we used a series of ten independent *gni2* triple-KO (aabbdd) mutants in the *cv.* Fielder background (Suppl Table S7); all lines showed the extended rachilla and increased floret number phenotype under greenhouse conditions (Figure 2C). It is worth noting that Fielder naturally carries the functional *GNI-A1105N* allele, indicating that the obtained *gni2* mutant phenotypes solely reflect triple-KO (aabbdd) effects. We therefore cultivated and randomized these T3- and T4-lines in soil, in small ∼1m^2^ plots with ∼300 plants per m^2^ under semi-field-like conditions in an S1-regulated aerated greenhouse together with small plots for Fielder (Figure 5A-C; Suppl Figure S4). We then hand-harvested the plots and threshed the spikes. To have a better estimate of the small-grain fraction of all lines, we also sieved the grains and subsequently obtained two grain fractions (only in GH171), i.e., larger (>2 mm) and smaller than <2 mm. Overall, we noticed a grain yield advantage in *gni2* mutant plots of approx. 10% when averaged across all grain fractions from all three environments (Figure 5D; Suppl Tables S8). We found that grain number and thousand grain weight responded inversely while harvest index (HI) was significantly improved in *gni2* mutants (Figure 5E). Notably, spike-bearing tiller number, vegetative biomass, and above ground weight remained unaltered (Figure 5F; Suppl Tables S9-S11), suggesting that the enhanced HI is likely reflects a shift in carbon allocations toward floral structures. Taken together, the superior performance of *gni2* mutant plants under high-density (∼300 plants/m^2^) and semi-field-like conditions (soil) underscores the value of these high-yielding KO-alleles at all three loci, including the previously unexplored *DD* genome.

**Figure 5.**
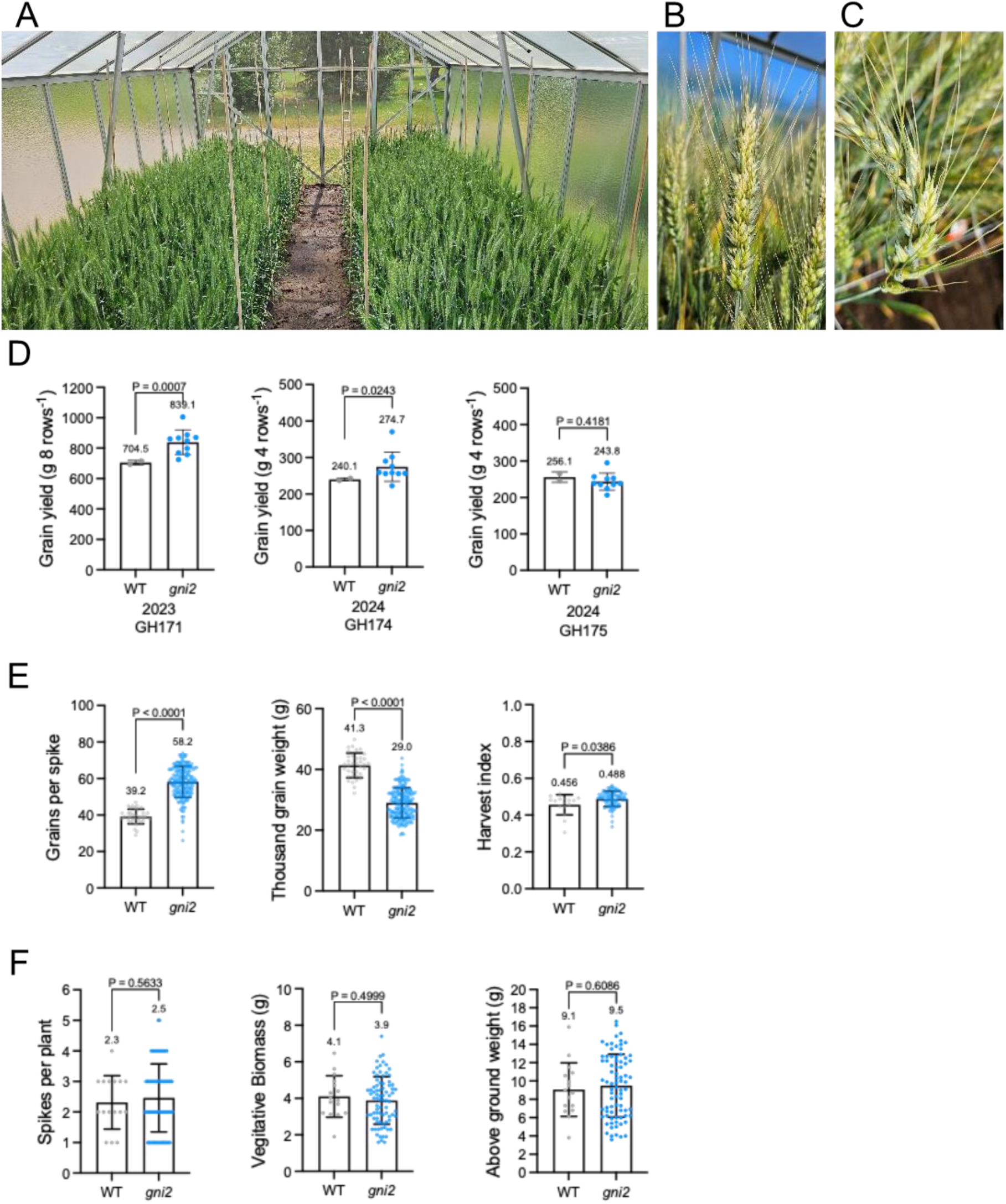
Grain yield and phenotypic effects of the loss-of-function triple *gni2* mutants in bread wheat. (A) Plants were soil-grown in contained S1-regulated greenhouses and a randomized design under semi-field-like conditions with a planting density of ∼300 plants per m^2^. (B) and (C) Representative spikes of the *gni2* triple mutants. (D-F) Phenotypic assessment of the *gni2* triple mutants in T3 and T4 generation of small plot-grown plants. Fielder is wildtype. All other CRISPR-edited lines of Fielder harbor triple-knockout mutations at the *GNI2* homoeo-loci. All plots were hand-harvested to obtain all grains. (E) Key component traits comprising grain yield, (F) spike-bearing tillers, vegetative biomass, and total above ground weight were determined. P-values were determined using the Welch’s t- test.

### *GNI2s* act as growth modulators, which work additively in root and shoot tissues, while concurrently extenuate distal floral growth via PTD-related transcriptional signatures

To seek a deeper functional understanding of *GNI2* actions, we queried public databases of hexaploid wheat for mRNA expression levels across various tissues and stages (Wheat eFP Browser; http://bar.utoronto.ca/efp_wheat/cgi-bin/efpWeb.cgi). Unlike *GNI-A1*, which is predominantly expressed in the developing spike and spikelets^9^ including floral organs such as ovary, stigma, and anther (Suppl Figure S5), all *GNI2s* are generally three to four times higher expressed and show the highest expression preference in roots and shoots during the entire life cycle of the wheat plant (seedling to milk stage), including some lower expression levels in floral tissues (Suppl Figure S6). To obtain more highly spatially resolved mRNA expression data from floral tissues, we sampled spikelets from tetraploid wheat Svevo around green anther stage. Within spikelets, we separated glumes and individual florets (floret 1 to floret 3), while floret 4 and the remaining apical floral structures were pooled into one sample (F4+n; Figure 6A), and applied gene expression analyses. We found that both *GNI2s* are predominantly expressed in F4+n samples thereby following a similar expression pattern as *GNI-A1* but to a slightly higher extent (Figure 6B), suggesting a similar functional growth modulation for *GNI-A1* and all *GNI2s* in apical florets of wheat spikelets. To better explore the transcriptional signatures and regulations during apical floret development and degeneration, we pursued an RNA-seq experiment of F4+n tissues from soil-grown greenhouse plants (Figure 6C-D) between wt Fielder and one triple KO mutant #30-3-11 (hereafter #30 or *gni2*) at two stages (tipping, TP; heading, HD). Our RNA-seq, GO-term, DEG, and hormonal analyses revealed that samples were clearly separated due to differential tissue fates (Figure 6D-E; Suppl Figure S7). While mutant #30 plants showed clear expression signatures for tissue growth, photosynthesis, cell division, and floral progression in their apical floral structures, the Fielder transcripts were strongly associated with senescence-, aging-, and stress-related processes (Figure 6D; Suppl Table S12), thereby indicating significant overlaps with a PTD (pre-anthesis tip degeneration)-related course of events in barley^23^. Very similar to floral PTD in barley, also floral PTD in Fielder wheat showed an enrichment for specific transcription factor families, such as MYB, bZIP, NAC, HD-Zip (*TaGT1*), while ABA-related transcripts (Figure 6D; e.g. cluster 4) and ABA levels were likewise upregulated (Figure 6E) accompanying the observed inhibitory effects of floral growth. Comparative analyses between wheat and barley data sets suggests that very similar transcriptional regulations determine apical floral fate in both related species.

**Figure 6.**
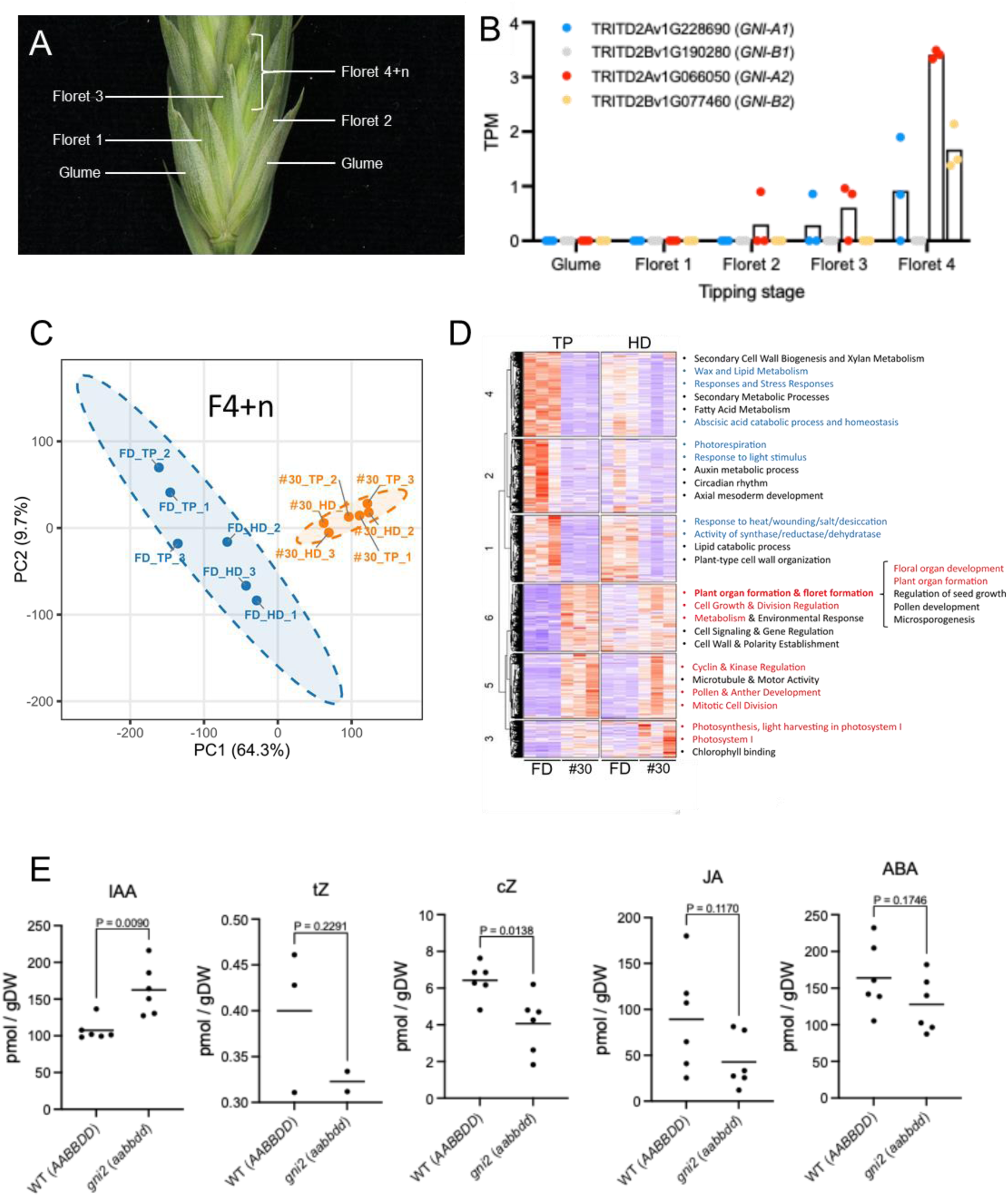
Transcriptional and hormonal analyses of wheat *GNI2.* (A) Spikelet of tetraploid wheat Svevo, including floral structures used for sampling in B. (B) Floral tissue expression patterns of *GNU* and *GNI2* in tetraploid wild-type wheat Svevo. (C) Principal component analysis using floret 4+n (F4+n) RNA-seq samples from wild-type hexaploid wheat Fielder (FD) and the *gni2* triple mutant (#30) at two stages (tipping, TP; heading HD). (D) Differential gene and GO term analysis of F4+n RNA-seq samples from wild-type FD and mutant (#30) at two stages. GO terms predominantly associated with FD are shown in blue (clusters 1, 2, 4) and largely related to stress responses. GO terms associated with mutant #30 are shown in red (clusters 3, 5, 6) and related to growth, organ formation, and floral greening. (E) Phytohormone profiling between wild-type FD and *gni2* triple mutant using floret 4+n at the tipping stage. IAA; indole-3-acetic acid, tZ; frans-zeatin, cZ; c/s-zeatin, JA; Jasmonic acid, ABA; abscisic acid. P-values were determined using the Welch’s t-test.

Since *GNI2s* are most highly expressed in roots during the three leaf-stage (Suppl Figure S6), we were equally inclined to explore how *GNI2s* affect root performance. To this end, we grew Fielder and *gni2* mutants in a rolled filter paper assay and measured in total 33 traits of eight days old seedlings, including shoot and root biomass, root length, root diameter, and root branching. Notably, none of the traits showed a significant difference between the wt and any mutant combination (Suppl Table S13), indicating that at least during early root growth and development both remained indistinguishable. These findings appear to be in line with an improved harvest index at maturity (Figure 5D), thereby supporting the idea of reproductive allocation in mutant plants towards floral structures, which in turn are less growth-inhibited, i.e. have lowered PTD for enhanced grain set.

## Discussion

Here, we uncovered a set of four homoeo-allelic haplotypes of *GNI2* in wheat. When paired with *gni-A1105Y* in bread wheat they enhance grain yield by about 5-7% under high-input farming and by 10-15% under low-input conditions, highlighting their potential to boost productivity across diverse farming environments. Our approach of merging genetic data with comprehensive wheat panels, which include multi-year, multi-environment grain yield data and various phenotypes, creates a strong and reliable basis for our association and grain yield studies. The observed lack of *DD* genome diversity for *GNI-D1* and *GNI-D2* even provides more future opportunities for improving grain yields in bread wheat; but only if matched with source-strong genotypes that are well adapted to high-density planting and exhibit high radiation use-efficiency^11,33–36^. This important pre-condition for future improvements largely stems from our observation that plants of the triple mutant *gni2* produced approx. 5 times more small grains (<2 mm) than Fielder (Suppl Tables S8). Therefore, the successful future utilization of these beneficial haplotypes requires assembling source-strong genotypes to express their full potential to better capitalize on the small-grain fraction. An alternative or complementary approach to overcome the small-grain-problem could be the post-anthesis application of affordable, environmentally friendly grain size enhancers like DMNB-T6P^37^ or brazide^38^ that may aide fill grains better.

Unlike in bread wheat, where natural *DD* genome diversity for *GNI1 and GNI2* remains untapped, all possible and important *loss-of-function* alleles for *GNI1* and *GNI2* in durum wheats have been identified. While allele frequencies for *gni-A1105Y* (∼95%) and *gni-B2* (45%) are high, the beneficial deletion-allele *gni-A2del* is only found in ∼5 to 19% of the accessions yet its origin could be traced back to important cultivars with prime spike fertility. For the future success of durum wheats, it will become equally important to unlock its full potential by using robust, source-strong genotypes that are locally adapted to perform at their best.

Although we did not observe the high-fertility floral phenotype (as reported in AltarC84 and others) in our bread wheat panels, we still identified significant allele effects from single *AA* and *BB* genome alleles that improved grain yield and its component traits. This suggests that, even without a clearly visible floral phenotype, these *GNI2* alleles can gradually influence other important traits—such as harvest index—through improved allocation mechanisms, ultimately contributing to increased grain yield.

Our analyses support a model in which *GNI2s* act as growth modulators that work additively in root and shoot tissues to alter reproductive allocation and harvest index. In the wheat spikelet, this is accomplished through local growth arrest of apical florets via PTD-related processes, indicating that spike PTD in barley and spikelet PTD in wheat share common features during the transcriptional reprogramming of slowly degenerating apical floral tissues. However, how *GNI2s* function during later root development and stem elongation to alter harvest index and reproductive allocation remains to be shown. Nonetheless, our study provides clear evidence that exploring PTD in temperate cereals in greater detail is a promising path for improving floret fertility and grain yield sustainably, even more so if it could be combined with the biological nitrification inhibition (BNI) trait for environment-friendly sustainable production^39^.

## Acknowledgments

We gratefully acknowledge M. Ando, A. Püschel, C. Trautewig, E. Weiss and K. Wolf for excellent technical support, Hannah Schneider and Ricardo F.H. Giehl for their valuable support while using the root scanner, all members from the PBP group for fruitful discussions, and support from IPK infrastructures.

## Accession numbers

The RNA-seq data have been submitted to the European Nucleotide Archive under accession number PRJEB108998.

## Materials and Methods

### Genotyping of the NCCR population

A balanced four-way multi-parental cross population, hereafter NCCR (Neodur, Claudio, Colosseo, and Rascon/2*Tarro) was used^40^. The total number of 334 NCCR lines were genotyped using a wheat-dedicated 90K Illumina Infinium array, including 81,587 effective SNPs^41^, and the linkage map was constructed (Milner et al. 2016).

### Phenotyping of the NCCR population

The phenotypic evaluation of NCCR population was carried out during two harvesting years (2010-2011 and 2011-2012) in three locations in the Po Valley: Cadriano (44°33’N lat., 11°24’E long. 2010-2011 and 2011-2012. Sizes of plots were, respectively, 2.4 m^2^ and 2.28 m^2^); Poggio Renatico (44°45’57” N, 11°25’31” E, 2010-2011. Plots: 4 m^2^) and Argelato (44°39’03” N, 11°20’34” E, 2011-2012. Plots: 4 m^2^). In each environment, as control genotypes, we inserted three replicates of the cultivars Levante, Meridiano, Orobel, Saragolla, and Svevo, as well as the four parental lines replicated twice. The 334 NCCR lines, the four parents, and the five control genotypes were evaluated according to a 19 x 19 α-lattice design^42^ with two replications, considered for each environment blocking and designed by means of FACTEX procedure (SAS Institute, 2006). Yield-related traits, such as spike length, number of fertile spikelets, number of unfertile spikelets, number of total florets per central spikelet, and number of grains per central spikelet. The central spikelet was considered as the 9*^th^* or 10*^th^*starting from the bottom of the spike; the bract (sterile floret) at the top of the spikelet was not included in floret counting.

### Genotyping of the Relief x Iride population

Relief x Iride biparental population was developed by Florimond Desprez (France). Relief is a French durum variety characterized by late flowering. Iride is related to AltarC84 with a short photoperiod and is highly productive. The population (1,550 genotypes), collected at the F4 generation, was multiplied in the Cadriano (UNIBO) field station (44°330N lat., 11°240E long., Bologna, Italy), during 2017/2018 and 2018/2019 growing seasons in 0.5 x 1 m plots, reaching F5 and F6 generations. These plants were tested in the Cadriano during 2019/2020 and 2020/2021 growing seasons in 0.5x1 m plots. These field trials were conducted in a complete randomized block design with two replicates using common repeated controls, such as Relief, Iride, AltarC84, and Svevo. Yield-related traits were evaluated as described above.

### Genotyping of the Svevo x Zavitan population

A population of 131 recombinant inbred lines derived from a cross between Svevo and Zavitan was grown in a glasshouse containing natural loamy soil with two open, non-glassed sides, each with porous metal netting to enable air circulation. We used approximately 10–15 cm distance between plants within a 1m row and 20–25 cm between rows. We used an incomplete block design with four single-plant replications arranged in 60 blocks. RILs were harvested from a greenhouse experiment, and the grain number per spike was recorded from the main culm. The number of grains per spikelet was calculated by dividing the number of grains per spike by the spikelet number. The Svevo x Zavitan RILs were genotyped using the 90K iSelect SNP array^43^, and the linkage map was constructed using 472 high-quality loci evenly distributed across the chromosomes^44^.

### Field data analysis

Data analysis was computed using statistical software R (R Core Team 2020) and SAS (SAS Institute Inc., 2006). ANOVA analysis was performed to calculate the effect of environmental variables (field trial effect and years), obtaining the variance component of each variable. Heritability for each trait was estimated using the heritability R package, as: *h*^2^ = *σ* 2g / *σ* 2g + (*σ* 2error/r). The phenotypic distributions that didn’t follow a normal distribution trend were normalized using the R package *BestNormalize*^45^. Best linear unbiased estimates (BLUEs) for each phenotypic trait were calculated on phenotypic data, including environmental variables as covariates, such as rows, columns, replicates, assessed number of plants, and the interaction between number of plants within the genotype. The Svevo x Zavitan RILs phenotypic traits were used to calculate BLUEs used for QTL analysis, and were calculated using the two-step mixed-model analysis implemented in Genstat (VSN International, Hemel Hempstead, UK). In the first step, we fitted the genotype and the replication as random effects with block nested within replication to determine the variance components used in the second step, with the genotype as a fixed term and the replication and nested blocks as random.

### QTL mapping

Composite interval mapping was conducted to detect QTL using mpMap, an R-coded platform specific for analyzing multi-parental populations^46^. Accordingly, marker-trait statistical associations are evaluated accommodating the conditional founder haplotype probabilities at each locus, given the available marker genotype data. Therefore, mpMap CIM QTL analysis will be further denoted as IBD-CIM. With mpMap, we performed interval mapping using the mpIM function (program ‘qtl’) using Simple interval mapping (SIM) and composite interval mapping (CIM) alghorithm^47,48^, fitting in ASReml v3.0^49^ a linear model and separately estimating fixed effects for each of the four founders along the chromosomes (Rascon/2*Tarro has been arbitrarily chosen as the reference haplotype). After Bonferroni’s correction, a *p−value* threshold of 0.05 was applied to each founder’s significance. We finally calculated the LOD-2 supporting interval (for combined data) and the LOD-1 supporting interval (for single-environment data) based on transformed *p −value* [*log*10(*p*)] profiles. The Svevo x Zavitan genome-wide scan of QTL was conducted using simple interval mapping (SIM). Then, the identified QTL were repeatedly used as co-factors in a composite interval mapping (CIM)^50^ until the QTL profile reached stability. The final QTL model following backward selection was fitted using REML and consisted of the overall mean (µ), the effect of each QTL (fixed), and the unexplained genetic and environmental residual (e, random), as implemented in Genstat and described below: Trait=μ+∑QTL+ e. The genetic interval included in the QTLs was explored using the Biomart tool included in the Ensembl plant database <u>(Durinck et al., 2009)</u>, where high-confidence genes were downloaded.

### Phenotyping of the BRIWECS panel

The phenotypic data of the BRIWECS panel of 191 bread wheat cultivars for 10 agronomic traits, which represent adjusted means over six locations and two growing seasons, were obtained from a previously published work^10^. The wheat cultivars were analyzed for grain yield [dt/ha], biomass [t/h], thousand kernel weight [g], grains per spike, grains per m^2^, spikes per m^2^, harvest index, plant height [cm], crude protein [%], protein yield [kg/ha].

### ONT sequencing of *GNI2* genes in 96 bread wheat cultivars from the BRIWECS panel

Genomic DNA for the BRIWECS wheat cultivars was extracted from young leaf tissues using the BioSprint 96 DNA Plant kit (Qiagen, Düsseldorf, Germany) according to the manufacturer’s recommendations. DNA concentrations were quantified using the Qubit dsDNA BR Assay kit from Invitrogen and a microplate reader with fluorescence excitation/emission (TECAN infinite 200, Männedorf, Switzerland). Specific primes (Suppl Table S14) targeting the entire gene plus promoter sequence of three *GNI2* homoeo-loci were developed based upon a multiple alignment of reference sequences previously published (including quality pseudomolecule assemblies and scaffolded assemblies published by the 10+ genome project). The first-round PCR amplification was performed in 25 µl containing 7 µl RNase-free water, 12.5 µl of GoTaq Long PCR Master Mix (Promega, Madison, WI, USA), 1.5 µl of each primer (10 µM), and 2.5 µl (40 ng/μl) genomic DNA. PCR reactions were performed in a T100 Thermal Cycler (Bio-Rad Laboratories, Hercules, CA, USA) using the following program: initial denaturing at 94 °C for 2 mins, followed by 35 cycles of 94 °C for 25 secs, 64.5°C annealing /extension for 6 mins up to 8 mins, with a final extension step at 72 °C for 10 mins. The amplified PCR products were checked by agarose gel electrophoresis. Afterwards, DNA quantity was measured, and equal amounts of PCR products of each gene for each cultivar were pooled into one tube and purified via AMPure XP beads (Beckman Coulter, Brea, CA, USA) to remove salts, primers, and proteins. The second round of PCR amplification, which aims to add barcode sequences into the amplicons, was carried out in 50 µl reaction volumes consisting of 25 µl GoTaq Long PCR Master Mix, 24 µl of the first-round PCR products, and 1 µl of a barcode primer (EXP-PBC096, ONT, Oxford, UK). PCR conditions used for barcoding were as follows: an initial denaturing step at 95°C for 2 mins, followed by 18 cycles of denaturation for 15 secs at 94°C, annealing for 15 secs at 62°C, and extension for 8 mins and 10 secs at 65°C, the final extension step was carried out at 65°C for 10 mins. PCR barcoding was followed by purification of the amplicons using AMPure XP beads, and DNA quantity was measured and equal amounts of all 96 cultivars (barcodes) were pooled into a single sequencing library. The MinION library was produced using the Ligation Sequencing Kit 1D (SQK-LSK110, ONT Oxford, UK) according to the manufacturer’s recommendations. About 30 fmol (150 ng) of pooled library was loaded and sequenced on MinION R9.4.1 flow cell for approximately 24 hours, until no further sequencing reads could be collected. After a complete the run, the flow cell was washed with Flow Cell Wash Kit (EXP-WSH004, ONT, Oxford, UK), and was used again for resequencing of the same pooled library.

### Base calling, ONT data filtering and alignment

The Raw electrical signals “fast5 files” obtained by the MinION instrument were processed using the base-caller Guppy version version 6.3.7+532d626 with model dna_r9.4.1_450bps_hac.cfg^51^ in a virtual machine operating with Ubuntu 20.04.1 LTS with two NVIDIA Tesla 4 TU104GL (NVIDIA Corporation) Graphic Processor Units (GPU). The guppy_barcoder was used to demultiplex the basecalled reads, with the option detect_mid_strand_barcodes. Reads with Q-score lower than 8, length less than 4000 bp were filtered out by using the NanoFilt v.2.8.0 tool^52^. Filtered reads were aligned against the *GNI2* sequences of wheat Fielder and Julius references^53–55^ using the NGMLR long-read mapper version 0.2.7^56^, with setting min-identity 0.80. Subsequently, the alignment files in SAM format were converted to sorted BAM files, with only reads with a map quality score above 50 were kept.

### Consensus sequences generation, variants calling, and haplotype analysis of *GNI2* genes from ONT data in the BRIWECS panel

Consensus *GNI2* sequences were generated by using reference-guided assembly method where reads were mapped against the reference sequence and then used to construct a consensus sequence using Amplicon_sorter tool and Canu assembler v2.2^57,58^. Polymorphisms were called from multi-alignment of consensus sequences aligning to a reference sequence with considering only regions present in a reference sequence using the SNP-sites tool^59^. We filter out SNPs with an allele frequency of less than 3% and SNPs located in homopolymeric regions. Detected SNPs were also visually inspected in the Integrative Genomics Viewer^60^. Based on SNPs identification from ONT sequencing, multiple haplotypes of *GNI2* genes were detected in the BRIWECS panel (Suppl Table S6).

### KASP marker design from Illumina 90K SNP Chip array, ONT sequencing data from durum and bread wheat accessions and genotyping on durum wheat mapping populations

Polymorphic SNP markers within *GNI-A2* confidence region between Svevo and AltarC84 were converted to KASP markers. KASP primers A and B were designed on the specific SNP being variety specific, whilst the common C primers was designed to be specific for the A genome on *T. turgidum* cv. Svevo^31^ KASP primers specificity was tested on Relief, Iride, Svevo and AltarC84 durum genotypes. Once confirmed, genotyping was performed on durum panel 1^61^ and Relief x Iride RILs *F*6 using KASP master mix kit and protocol (LGC genomics). Relief x Iride genotypes were sown and harvested for DNA extraction using CTAB extraction protocol^62^. A and B primers were attached with different tails complementary to different fluorescent probes (FAM and HEX) included in the master mix, to discriminate different allelic variants on specific SNPs: GAAGGT- GACCAAGTTCATGCT (5’-FAM) for primer A and GAAGGTCGGAGTCAACGGATT (5’-HEX) for primer B. Primer C was common to both primer A and B and was designed on homoeologs SNP for genome A. Additional KASP markers were converted from Affimetrix 420K in order to fine map the *GNI-A2* confidence interval (Suppl Table S14) Furthermore, four KASP markers were developed from the polymorphisms detected by ONT to distinguish cultivars carrying different haplotype variants of *GNI2* in BRIWECS panel (Suppl Table S14). The PCR protocol was used according to the manufacturer’s instructions by LGC Genomics. The following thermal protocol was used: 94°C 10 minutes – 10 cycles touchdown at 94°C 10 seconds and starting from 65°C to 57°C, 26 cycles at 94°C for 20 seconds and 57°C for 60 seconds. Further re-cycling steps were applied in case a better separation of clusters was required using the following thermal protocol: 3 cycles at 94°C for 20 seconds and 57°C for 60 seconds.

### Haplotype analysis of *GNI2* homoeologs using the GABI panel

A panel of 197 GABI wheat cultivars^14^ was re-sequenced using the primers listed in Suppl Table S14. Phenotypes of the studied cultivars were recovered from archived data^14^.

### ONT sequencing analysis on *GNI-A2* and *GNI-B2* in the tetraploid wheat global collection

Ninety-six genotypes representing diverse tetraploid wheat^31^ were sequenced using specific primer pairs specifically designed for the *GNI-A2* (*TRITD2Av1G066050*) and *GNI-B2* (*TRITD2Bv1G077460*), covering 4 kbp upstream of the ATG, and downstream of the stop codon to include promoter and terminal regions (Suppl Table S14). PCR amplicons for each genotype were barcoded and sequenced. Reads were then filtered for stringent parameters, demultiplexing reads containing both right and left barcode corresponding to each genotype. Reads were mapped against *Triticum turgidum* cv. Svevo Platinum genome using *Minimap2* (Li et al., 2018). Assemblies were generated using three software: Amplicon SorteR^58^, FLYE^63^, and NECAR^64^. For each genotype, the assemblies obtained with the three software were compared, multi-aligned, and manually curated to select the most consistent one. The multi-alignment of 96 consensus for each amplicon was then manually inspected and curated, and SNPs were called using the msa2vcf.py software^65^ and SNPs’ effects were called using the ensemble VEP^66^ using the final Svevo Platinum annotation.

### Screening the tetraploid germplasm for the *GNI-B2* loss-of-function mutation

A CAPS marker targeting the 1bp deletion in the *GNI-B2* coding region was developed. PCR amplification (amplicon size 484bp) was conducted, followed by digestion with Hpy188III, specifically digesting the functional allele (Suppl Table S14). The *GNI-B2* indel marker was used to characterize the presence/absence of the indel in a panel of 290 tetraploid accessions (Suppl Table S3) selected from two previously studied diversity panels^67,68^.

### RNA-seq analysis

The clean RNA-Seq data were mapped to the Chinese Spring v1.1 reference genome^69^. Kallisto^70^ (v) was utilized to determine gene expression values (TPM) and counts with each annotated gene. DESeq2(v1.51.7)^71^ was used to identify differentially expressed genes (DEGs). (Add description of control group and experimental group) Genes meeting the criteria of log2 (fold change with count) > 1 and false discovery rate (FDR) < 0.05 were considered as DEGs. GO enrichment analyses for target genes were conducted using the topGO(v2.46.0) package in R (https://github.com/federicomarini/topGO).

### Generation of constructs with *cas9* and gRNAs

Twenty nucleotide (nt) target sequences present immediately adjacent to a Protospacer Adjacent Motif (PAM) were selected using DESKGEN^TM^ CRISPR Libraries and verified using the RNAfold web server (http://rna.tbi.univie.ac.at/cgi-bin/RNAWebSuite/RNAfold.cgi) according to Kumlehn et al.^72^ for the *GNI2* gene. Four single guide RNAs (gRNAs) for *GNI2* (target-specific parts of gRNA1: GCTGGAGCTGAGCTTCCGGG, gRNA2: GAGGGGATCCCAAGAAGCGG, gRNA3: GAGGGCGCGGCGCAGGAGG, and gRNA4: ACAGGGGGAGAGCCGGCCGA) were selected. The constructs targeting *GNI2* (four guides) were generated using the CasCADE modular vector system, which is based on a hierarchical Golden Gate cloning strategy (Hoffie et al., in prep). Oligos (Table SX) with complementary target sequences and overhangs for cloning into BsaI restriction sites were annealed and cloned into BsaI–linearized vectors pIK5 to pIK8 containing the *TaU6* promoter. The resultant four vectors containing four gRNA units were digested with Esp3I enzyme and assembled into one vector, pIK19 (four guides). To combine the multiple gRNA fragments with the *cas9* expression unit (pIK83) and an auxiliary unit vector (pIK155) into vector pIK22, BsaI restriction digest followed by ligation was used. Last, all expression units were mobilized into binary vector p6-d35S-TE9 (DNA Cloning Service) via SfiI restriction digest.

### Transformation

The spring wheat cultivar ‘Fielder’ was used for transformation. To obtain immature embryos, wheat plants were grown continuously. Plants were grown in a glasshouse with day/night temperatures of 16°C/10°C under a 10-h light/14-h dark photoperiod for 10–12 weeks. Just before the heading stage, the plants were transferred to a controlled environmental chamber with day/night temperatures of 20°C/13°C under a 14-h light (300–500 µmol/m2/s)/10-h dark photoperiod. Spikes were collected at 14–17 days after anthesis, and immature embryos were isolated aseptically. Inoculation with Agrobacterium (*Agrobacterium tumefaciens*) harboring the <vector name> plasmid and generation of transgenic wheat was performed as described previously^73^. In brief, the immature embryos were centrifuged, then inoculated with Agrobacterium. After 2 days of co-cultivation, the embryo axes were removed, and the remaining scutella were cultured without the selective agent, hygromycin B, for 5 days. To increase genome editing efficiency, the scutella were treated at 30°C for the first 1day of this non-selective culture step^74^. After the selection for hygromycin resistance, proliferated calli were transferred to the regeneration medium. Regenerated plants were transferred to the rooting medium, and then rooted T0 plants were transplanted into the soil. The transgene integration in T0 plants was detected by genome PCR using the following primers: HPT_F: 5’- GTGTCACGTTGCAAGACCTG-3’ HPT_R: 5’-GATGTTGGCGACCTCGTATT-3’ The transgene-integrated plants were grown at 20°C/13°C under a 14-h light/10-h dark photoperiod in a controlled environmental chamber.

### Measurement of plant hormones

Wheat florets at the tipping stage were harvested, their fresh weights measured and then frozen in liquid nitrogen. Frozen florets were crushed into a fine powder with a zirconia bead (diameter 5 mm). Stable isotope-labeled internal standards were added to the samples at the start of extraction, and plant hormones were extracted, purified, and separated into three fractions as previously described^75^ without MS probe derivatization. Abscisic acid (ABA), auxins, gibberellins, jasmonic acid (JA), salicylic acid (SA), and cytokinins (CKs) were analyzed by ultra-performance liquid chromatography (UPLC) on an octadecyl-silica (ODS) column (ACQUITY Premier HSS T3 with VanGuard FIT, 1.8 µm, 2.1 × 100 mm; Waters, Milford, MA, USA) coupled to a tandem quadrupole mass spectrometer (ACQUITY UPLC System/Xevo TQ-XS; Waters) equipped with an electrospray ionization (ESI) source. Each fraction was analyzed in a separate run under optimized LC-MS/MS conditions.

### Phenotyping of roots and seedlings

#### Experimental Design

The experiment comprised eight hexaploid wheat (*Triticum aestivum*) genotypes carrying all combinations of wild-type and loss-of-function *GNI2* alleles across the *AA*, *BB*, and *DD* genomes: one wild-type control (*AABBDD*), three single mutants, three double mutants, and one triple mutant. Three filter papers served as blocks, with three to four biological replicates per genotype, resulting in a total of 80 experimental units.

#### Germination and Growth Conditions

Grains were germinated, and seedlings were transferred to 198 × 592 mm filter papers. Three to four grains were placed along a 5 cm reference line; the papers were stacked and rolled into scrolls, and each scroll was placed upright in a 5-L container containing 1.5 L of Hoagland nutrient solution. Containers were lined with filter paper to maintain moisture and covered with aluminum foil to restrict light exposure to the root zone. Seedlings were subsequently transferred to greenhouse conditions, and the nutrient solution was replenished twice during the experiment. After two weeks, roots were imaged using a water-immersion setup with an Epson Expression 12000XL flatbed scanner. Shoot and root fresh weights were recorded immediately, and dry weights were obtained after one week of drying.

#### Trait Measurements and Root Image Analysis

Root system architecture was quantified from scanned images using RhizoVision Explorer (version 2.0.3)^76^. Images were acquired at 600 dpi (0.0423 mm pixel⁻¹), processed using a fixed global threshold (231), and the largest connected component was retained. Background noise was removed by filtering components ≤1 pixel, and root pruning was applied at a threshold of 5 pixels. In total, 33 traits were recorded, including shoot and root fresh and dry weights, total biomass, water content, root-to-shoot ratio, root architecture parameters (total length, number of tips, number of branch points, branching frequency, network area, and perimeter), root diameter descriptors, and diameter class–based metrics for length, projected area, surface area, and volume across three classes (0-0.2, 0.2-0.3, and >0.3 mm; Suppl Table S13).

#### Statistical Analysis

All statistical analyses were performed in R (version 4.5.3). For each trait, normality and homoscedasticity were assessed using the Shapiro-Wilk and Levene’s tests, respectively. Traits meeting both assumptions were analyzed using linear models including genotype, filter paper, and replication, followed by Dunnett’s test against the wild type (*AABBDD*). Traits with heterogeneous variances were analyzed using Welch’s ANOVA followed by Games-Howell post hoc tests. Traits that did not meet normality assumptions were analyzed using the Kruskal-Wallis test, followed by pairwise Wilcoxon rank-sum tests with Holm correction. Significance was defined at *p* ≤ 0.05. A full list of traits and tests is provided in (Suppl Table S13).

---

**Supplementary Figure S1.**
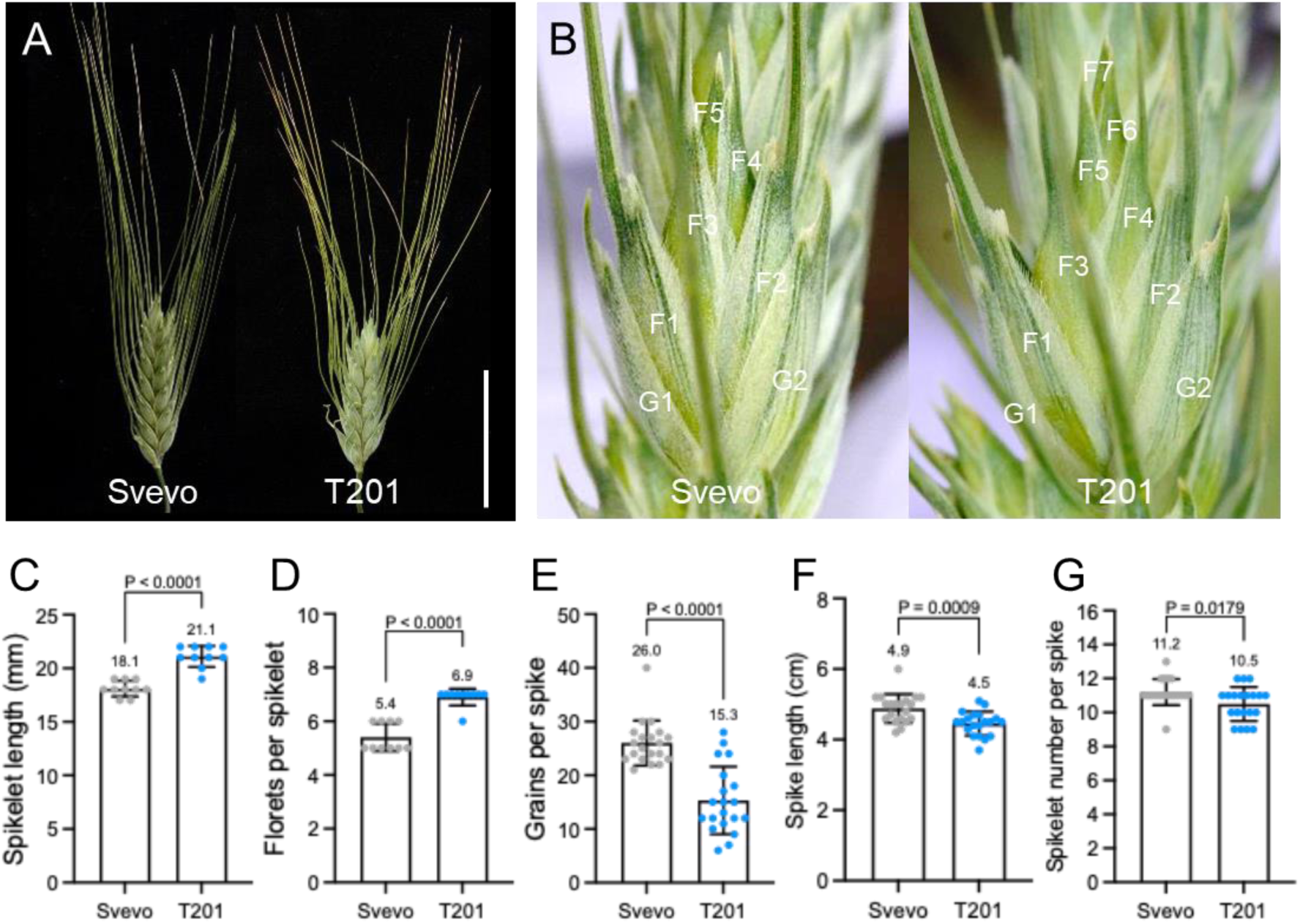
Phenotypic characterization of the *gni-A1_105Y_lgni2* triple mutant in tetraploid wheat. T201 was derived from an EMS mutagenesis TILLING-screen of *cv.* Svevo. T201 possesses naturally the two in durum wheat predominantly occurring *gni-A1_105Y_* (reduced function) and *gni-B2_29S l)e_i* (loss-of-function) alleles, but also the EMS-induced *gni-A2_L118F_* substitution. (A) Comparison of spike phenotypes of Svevo and T201. (B) Representative spikelet comparison with more florets per spikelet in the mutant T201. (C-G) Phenotypic assessment of spike traits. P-values were determined using the Welch’s t-test.

**Suppl Figure S2.**
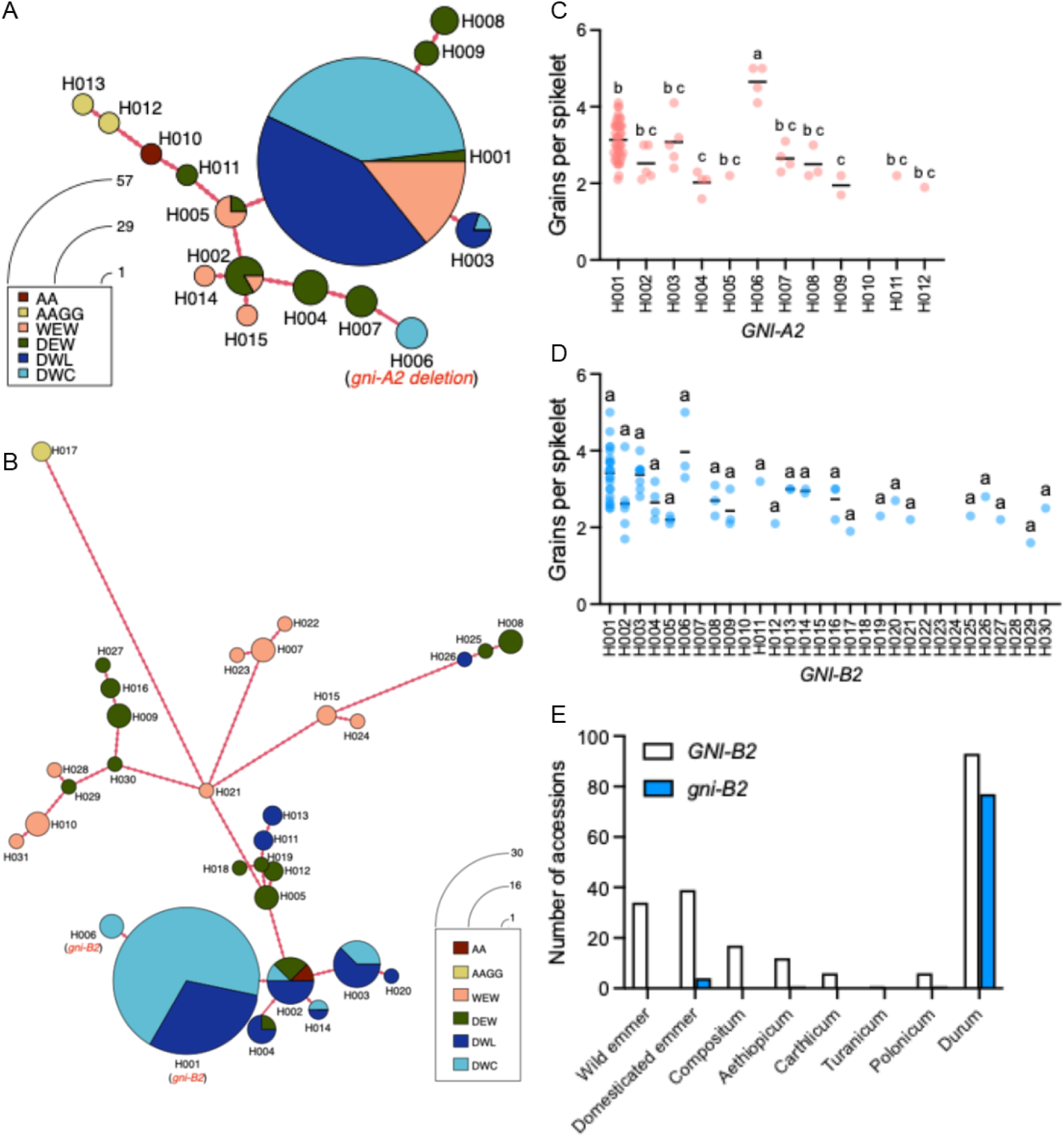
Allelic variation at *GNI2* in tetraploid wheat. (A) and (B) Haplotype networks in a panel of 96 tetraploid wheat global collection (TGC). (A) *GNI-A2* haplotypes. (B) *GNI-B2* haplotypes. AA: *Triticum urartu,* AAGG: Timopheevi, WEW: wild emmer wheat, DEW: domesticated emmer, DWL: durum wheat landraces, DWC: durum wheat cultivars. (C) *GNI-A2* haplotype 6 (H006) cultivars produce the highest number of grains per spikelet. (D) No significant changes were observed among *GNI-B2* haplotypes. Horizontal bars indicate the mean. (E) *GNI-B2* allele frequency in a panel of 290 wild and domesticated tetraploid accessions. The letters in (C) and (D) indicate where means differed from one another significantly *(P <* 0.05) as determined by Tukey’s honest significant difference test.

**Suppl Figure S3:**
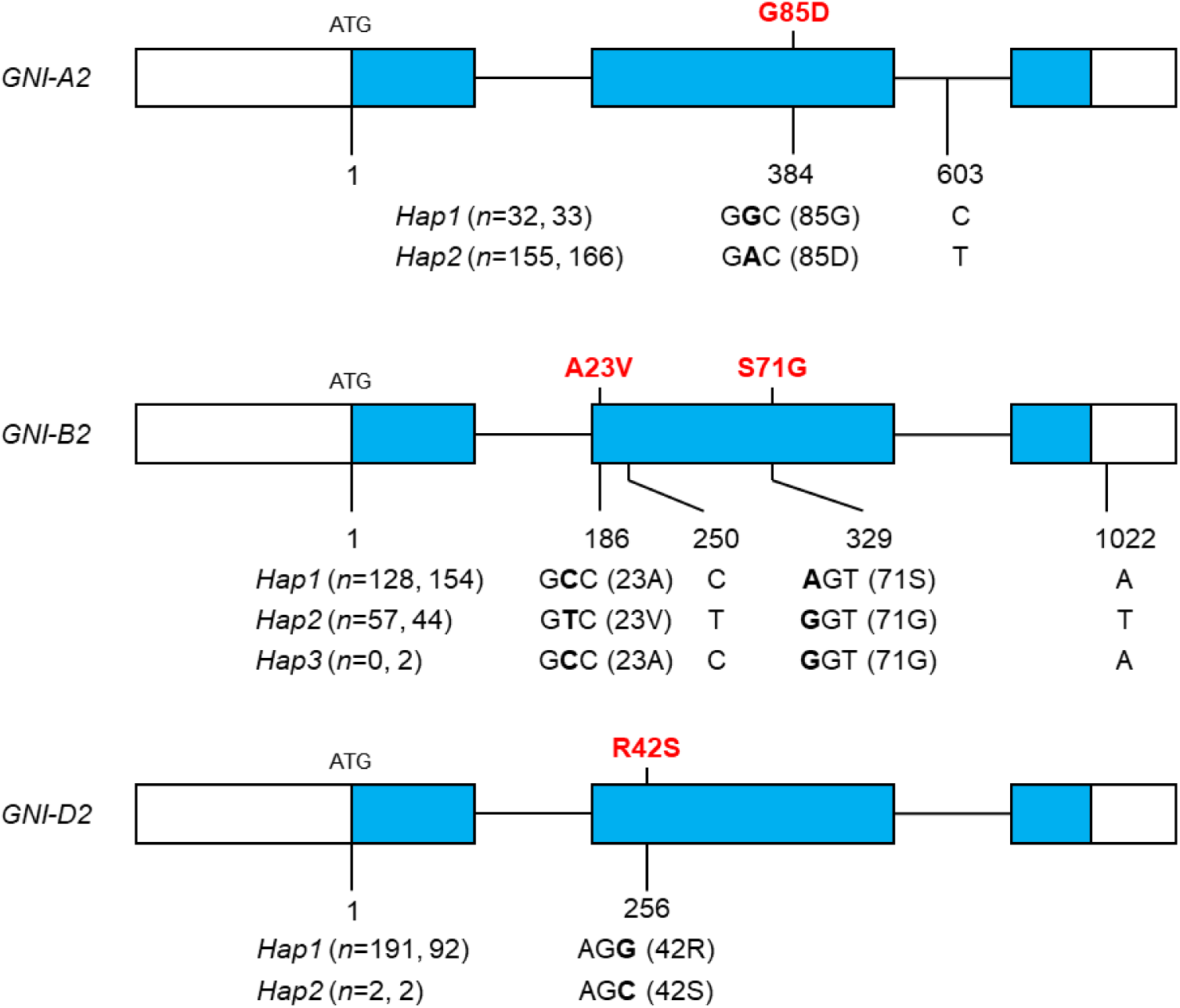
Allelic variation at *GNI2* in bread wheat. A panel of 210 European winter bread wheat cultivars (GABI) and panel of 199 bread wheat cultivars (BRIWECS) were re- sequenced.Hap 1 refers to the sequences from chines Spring. White boxes indicate untranslated regions, blue box coding sequences, and horizontal lines introns.

**Suppl Figure S4.**
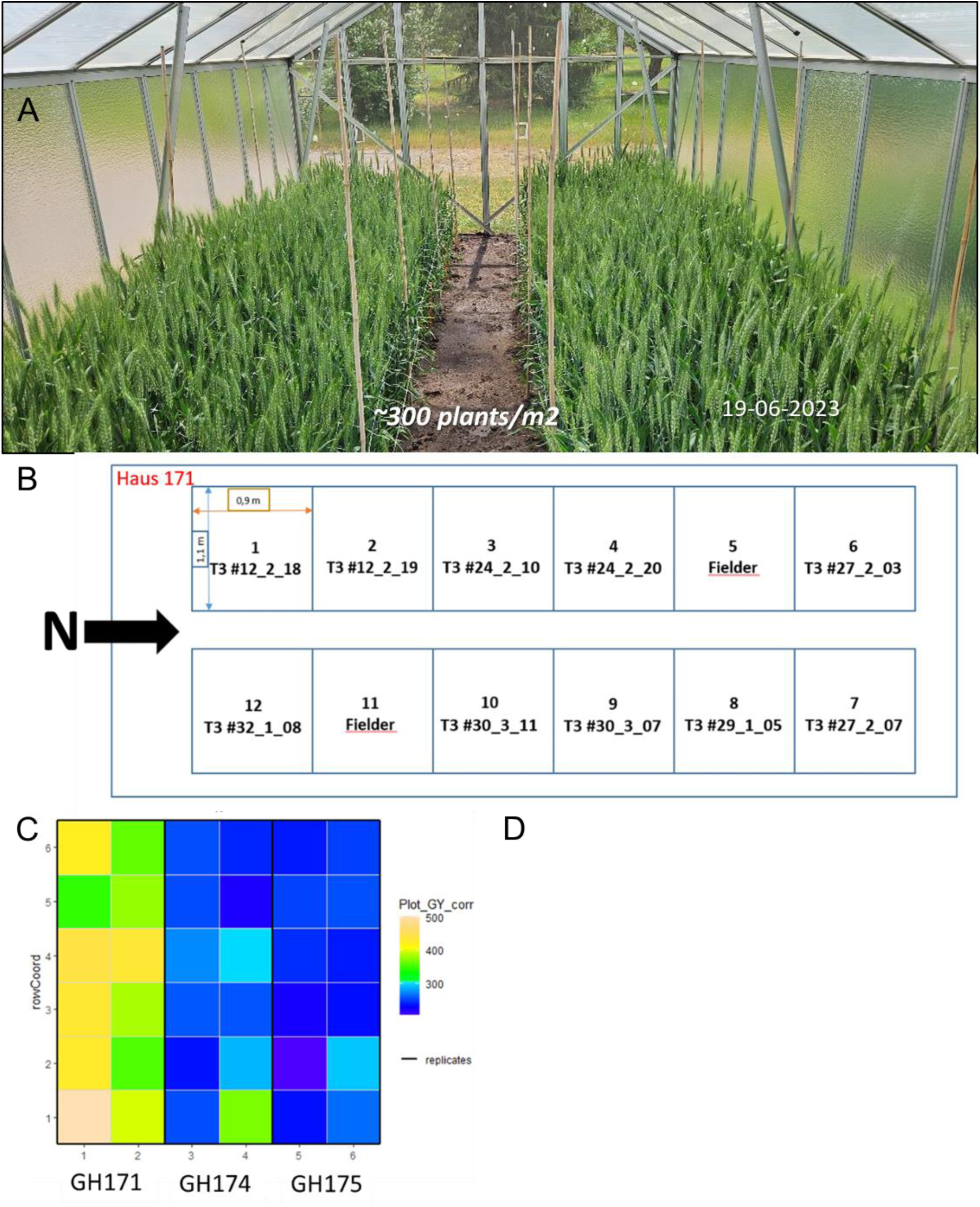
Effects of the loss-of-function mutations of *GNI2* in bread wheat. (A) Plants were soil-grown in contained S1 -regulated greenhouses under semi-field-like conditions with a planting density of-300 plants per m^2^ in 2023 (GH171) and 2024 (174,175). (B) Randomized design showing the different T3 plants in GH171 and Fielder in approx. 1 m^2^ plots. All phenotypic traits were derived by harvesting the core of the plots (∼1/4 m^2^). (C) Heat map of plot grain yields in the three greenhouses used for spatially adjusting mean values for the obtained phenotypic traits.

**Suppl Figure S5:**
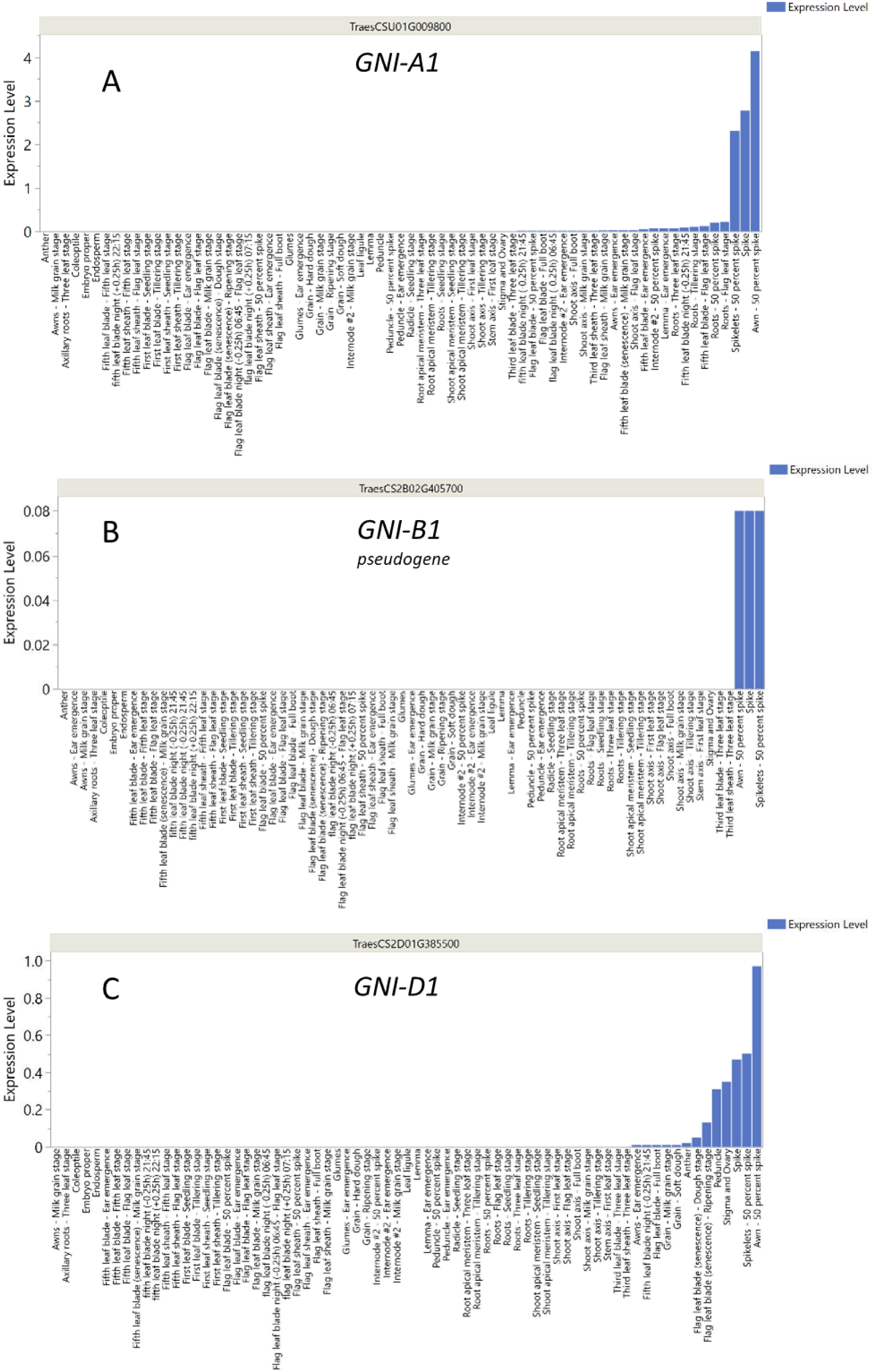
*GNI1* homoeologous mRNA expression levels from *cv*. Azhurnaya in hexaploid bread wheat. Shown are the mRNA levels from various tissues and stages (source: ??) in (A) for the AA genome, (B) the BB genome, and (C) the DD genome copy

**Suppl Figure S6:**
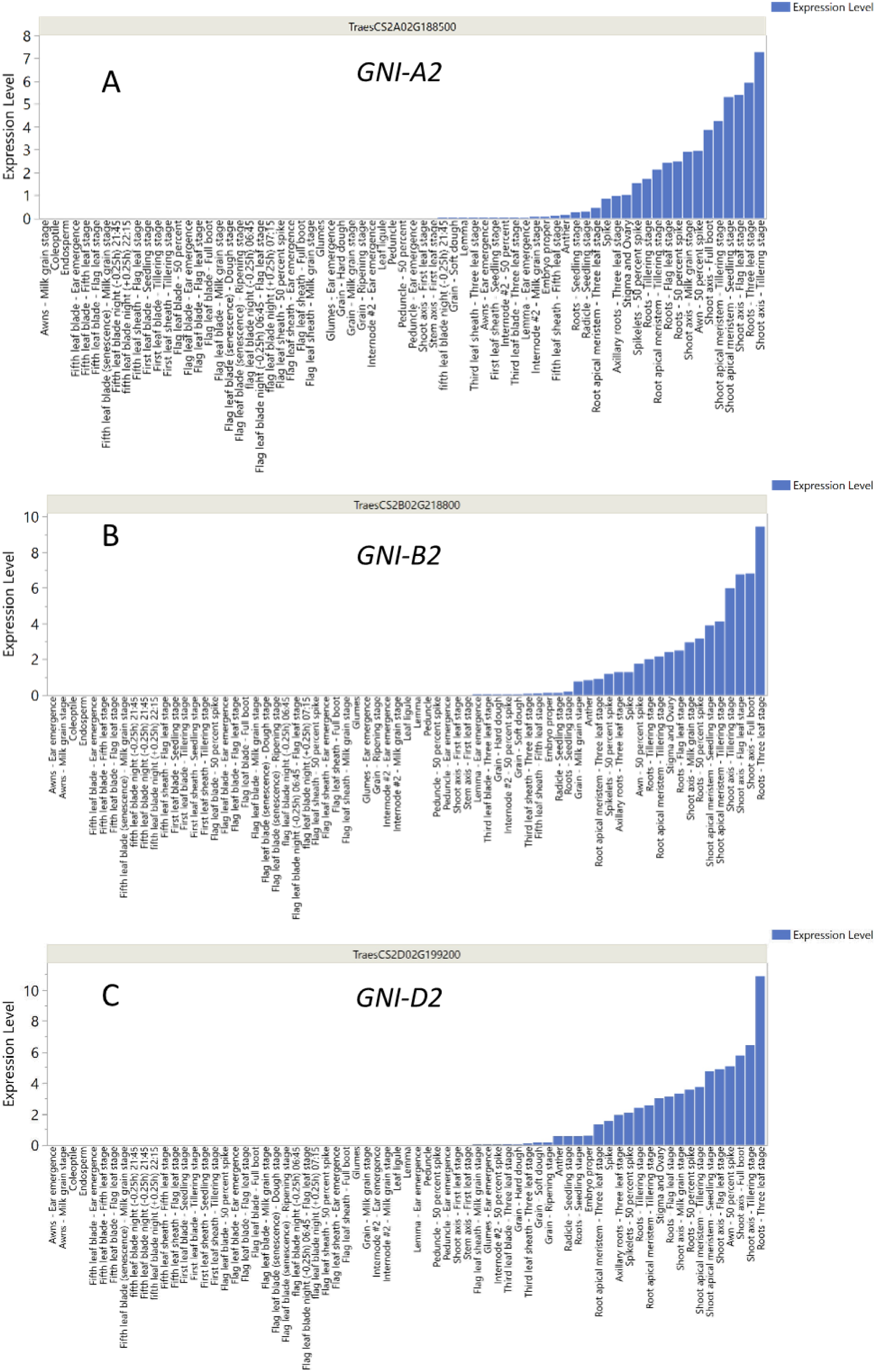
*GNI2* homoeolgous mRNA expression levels from cv Azhurnaya in hexaploid bread wheat. Shown are the mRNA levels from various tissues and stages (source:??) in (A) for the AA genome, (B) the genome, and (C) the DD genome copy

**Suppl Figure S7:**
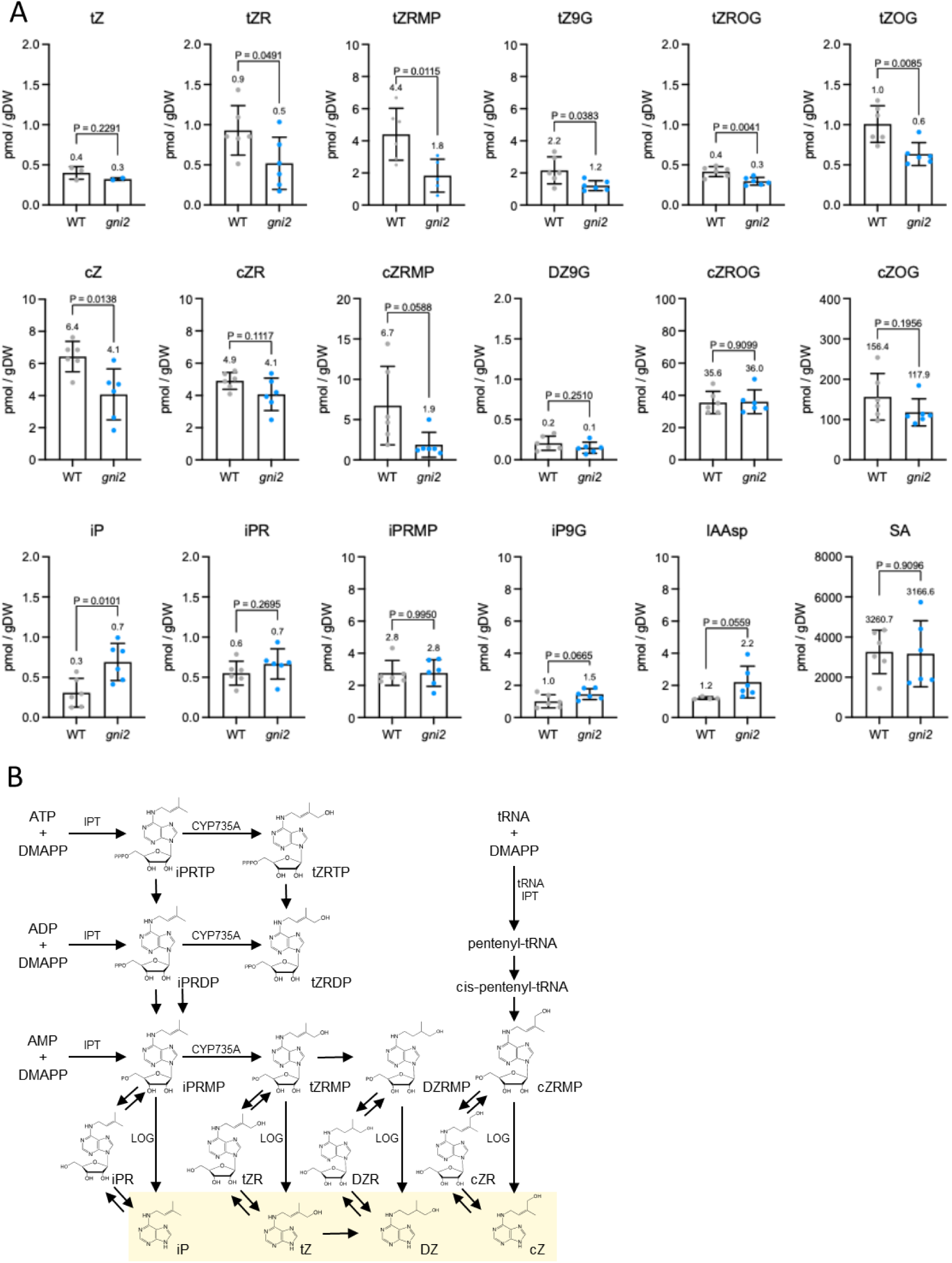
Comparison of phytohormones in wild-type Fielder and *gni2* triple-KO mutant. (A) Phytohormone profiling from upper florets at the tipping stage. P- values were determined using the Welch’s t test. (B) Cytokine biosynthetic metabolic pathway. IPT; adenosine phosphate isopentenyltransferase, CYP735A; Cytochrome P450 monooxygenase, LOG; LONELY GUY, iPRTP; iP-riboside 5’-triphosphate, iPRDP; iP- riboside 5’-diphosphate, iPRMP; iP-riboside 5’- monophosphate, iPR; iP riboside, iP; isopentenyl adenine, tZRTP; tZ-riboside 5’ triphosphate, tZRDP; tZ-riboside 5’- diphosphate, tZRMP; tZ-riboside 5’- monophosphate, tZR; tZ riboside, tZ; transzeatin, DZRTP; DZ-riboside 5’ triphosphate, DZRDP; tZ-riboside 5’-diphosphate, DZRMP; DZ-riboside 5’-monophosphate, DZR; DZ riboside, DZ; dihydrozeatin, cZRMP; cZ- riboside 5’-monophosphate, cZR; cZ riboside, cZ; cis-zeatin.

