## Supplementary material for "Homoeo-alleles of wheat *GNI2* fuel grain yields across input environments": Suppl Info

1. Leibniz Institute of Plant Genetics and Crop Plant Research (IPK), 06466 Seeland, OT Gatersleben, Germany
2. Faculty of Agriculture, Tottori University, Tottori 680-8553, Japan
3. Plant Genetics and Breeding Department of Agricultural and Food Sciences DISTAL, University of Bologna, 40127 Bologna, Italy
4. Department of Plant Breeding, Justus Liebig University Giessen, Giessen, Germany
5. Institute of Crop Science, NARO, Tsukuba, Japan
6. International Maize and Wheat Improvement Center (CIMMYT), Mexico City 06600, Mexico
7. RIKEN Center for Sustainable Resource Science, Yokohama 230-0045, Japan
8. Faculty of Natural Sciences III, Institute of Agricultural and Nutritional Sciences, Martin Luther University Halle-Wittenberg, 06120 Halle, Germany
9. Cluster of Excellence on Plant Sciences (CEPLAS) ‘SMART Plants for Tomorrow’s Needs’

^#^ Current address: The Robert H. Smith Faculty of Agriculture, Food & Environment, The Hebrew University of Jerusalem, Rehovot 7610001, Israel.

^§^ Current address: Heinrich-Heine-Universität Düsseldorf, Faculty of Mathematics and Natural Sciences, Centre for Plant Genome Engineering, Düsseldorf, Germany.

🖂**Corresponding authors:**

Marco Maccaferri

Thorsten Schnurbusch

**Plant materials used for QTL cloning**

To detect natural variants underlying floret fertility and increased grain numbers, we selected parental lines of our populations using AltarC84, a hallmark, high-fertility durum CIMMYT variety, Iride, and a four-way cross among Neodur-Claudio-Colosseo-Rascon/2*Tarro (NCCR, 300 genotypes), where Iride and Rascon/2*Tarro are both derivatives of AltarC84. For detecting *QGns.ipk-2B*, we used a biparental population obtained from Svevo x Zavitan, in which Svevo is a hallmark, fertile, high-quality Italian durum variety, and Zavitan is a wild emmer wheat from Israel. Both parents had high-quality reference genomes^1^, showing heritable genetic variation for grain yield components with a focus on the number of fertile florets and grains per spikelet or spike (**Main Text Figure 1**). We observed that Svevo showed higher floret fertility than wild emmer Zavitan with Svevo showing 49.6 grains per spike and 3.34 grains per spikelet, while Zavitan developed on average 29 grains per spike and 1.63 grains per spikelet.

Based on two years of field experiments in multiple environments, the NCCR mapping population showed a major QTL cluster for spike fertility traits on chr 2A, hereafter referred to as *QGns.ubo-2A* = *GNI-A2* in a wide, low recombinogenic interval of 8 cM, between the NCCR recombination bins located at 81.1 cM (IWB26960, corresponding to 83,047,998 Mb of Svevo v1 reference genome) and 89.1 cM (IWA581, 154,732,761 Mb) (**Suppl Data S1-2**). The most associated marker was IWA581, which had a phenotypic effect of 39.20% in the NCCR population and a -logP value of 42.22 (**Suppl Data S1-2**). For spike fertility, the *GNI-A2* locus was associated with an *R*^2^ of 39.2%, combined across years and environments, and with an additive effect relative to Neodur parent equal to -0.04 (ns) for Claudio, -0.01 (ns) for Colosseo, and + 0.54 (****) for Rascon/2*Tarro for grain number per spikelet. *GNI-A2* was clearly distinct from the *Ppd-A1* region, which mapped in the same population (Milner et al., 2016) to 36,577,899 bp of the Svevo genome corresponding to 55.6 cM or 114.8 Mb distal to *GNI-A2*. In the *GNI-A2* region, the NCCR polymorphic Illumina wheat 90K SNP showed the presence of two clearly distinct long-range haplotypes, one carried by the CIMMYT parent Rascon/2*Tarro and the other shared by the three Neodur, Colosseo, and Claudio parents (**Suppl Data S1**).

We fine-mapped *QGns.ubo-2A* = *GNI-A2* using recombinants identified in 1,550 individual F5 plants derived from a cross between *cvs*. Relief and Iride (**Main Text Figure 1C**) segregating grain number per spike. The *GNI-A2* interval was explored considering the chromosome region with the highest association scores in the NCCR population corresponding to the 149-156 Mb interval (**Suppl Data S1-3**), where 16 Illumina 90K wheat SNPs were converted to single fluorescent KASP® markers (Makhoul et al. 2020), evenly probing the *GNI-A2* confidence interval and suitable for screening novel recombinant lines, and fine-mapping the QTL and inspecting natural haplotype diversity. Twelve polymorphic KASP® markers (namely KUBO-47, KUBO-44, KUBO-177, KUBO-178, KUBO-179, KUBO-169, KUBO-182, KUBO-173, KUBO-175, KUBO-181, KUBO-49 and KUBO-50) were used to genotype Relief x Iride F_6_ population: 401 genotypes showed the WT (Relief) haplotype, 326 accessions had the *GNI-A2* positive allelic effect and (+/+) haplotype, and 10 recombinant genotypes were identified (**Suppl Data S3-4**). A summary of the steps performed for fine-mapping and GNS variability of mapping population parental/recombinant lines is depicted in **Suppl Info Figure 1A-B**. Based on the Svevo and Chinese Spring *GNI-A2* sequence information, including their homoeologs, we developed a *GNI-A2*_Deletion-specific PCR marker (**Suppl Figure 1C**). The deletion *gni-A2_del_* -specific PCR assay was validated on all the parental lines used in this study and in additional high floret fertility successful cultivars directly derived from AltarC84, such as the cultivars CIMMYT Atil_C2000, as well as the Italian Saragolla.

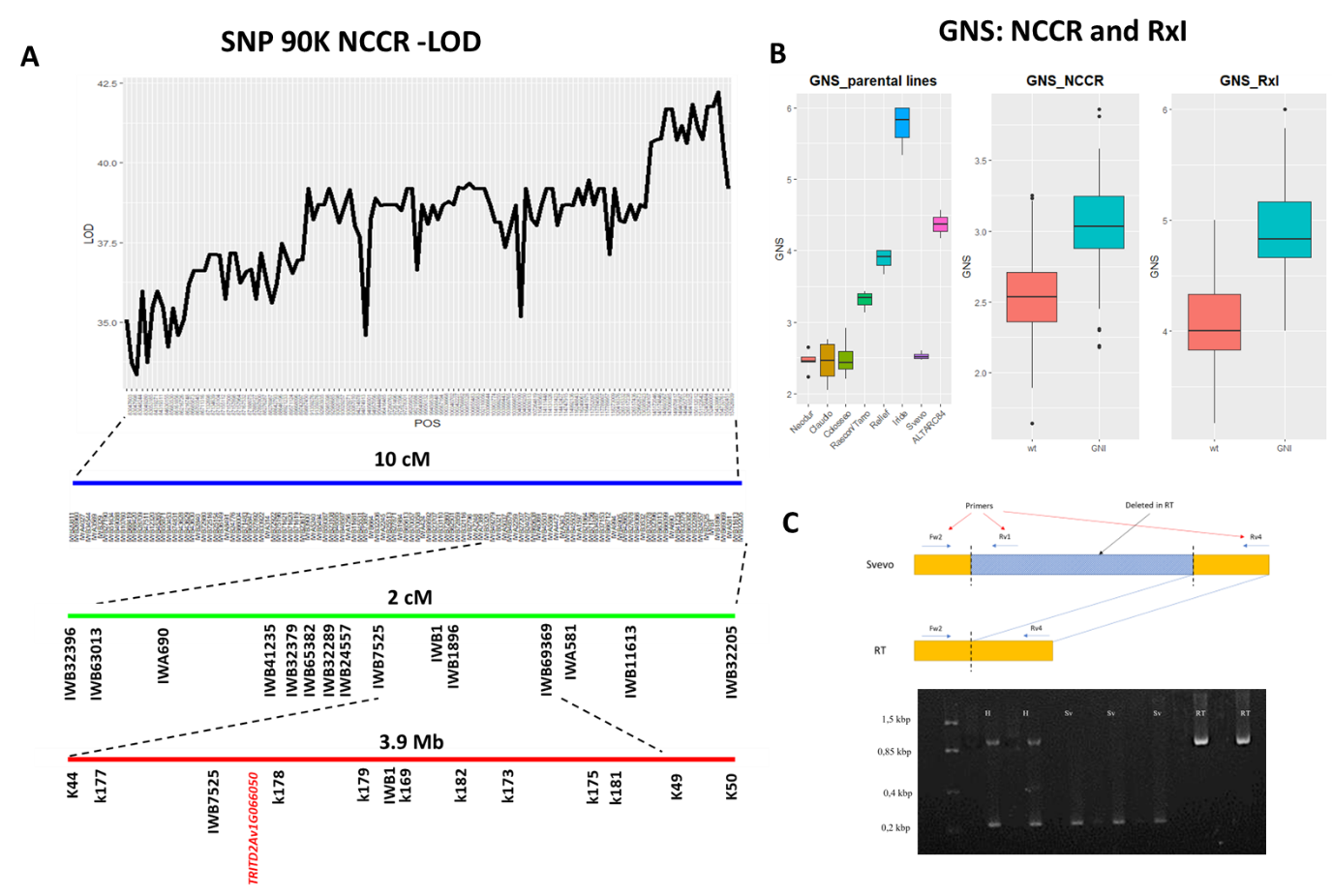

**Supp. Info Figure 1:** Map-based identification of *GNI-A2*. (A) Fine-mapping based on the NCCR multiparental and Relief x Iride (RxI) biparental populations. The genetic interval was delimited to 3.9 Mbp with 11 KASP® markers. (B) Variation of grain number per spikelet (GNS). (C) Validation of the *TRITD2Av1G066050* deletion. Sv: Svevo, H: heterozygote samples.

**Assessing nucleotide diversity at the *GNI2* homoeo-loci and the origin of the AltarC84 deletion allele**

The allele frequencies and haplotype diversity of the *GNI-A2* and *GNI-B2* were estimated in a panel of 96 tetraploid global collection (TGC)^2^. This includes *T*. *urartu* (AA), *T. timopheevi* (AAGG), wild emmer wheat (WEW), domesticated emmer wheat (DEW), *durum* wheat landraces (DWL), and durum wheat cultivars (DWC). The haplotype network of the *GNI-A2* interval identified 15 haplotypes, each one characterized by a different composition of the six main subpopulations (**Supp. Info Table 1**). We found that the complete AltarC84 Hap-A6, including the *GNI-A2* deletion, post-dates the ancestral emmer SNP-based haplotype, as none of the emmer accessions possessed the *GNI-A2* deletion, clearly suggesting post-domestication selection of this haplotype.

Hap-A1 corresponds to the wild type (Svevo) genotypes, and it is highly represented in DWC, DWL, WEW, and in a minor part of domesticated emmer wheat, with a frequency of 57 among all the 96 selected TGC genotypes. The *gni-A2_del_* deletion allele (Hap-A6) identified in AltarC84 had a highly significant positive effect for grains per spikelet in the same panel (**Suppl Figure S2A+C**) but at a very low frequency, i.e. present in 4 out of 96 accessions core and occurred only in known high-fertility cultivars, such as CIMMYT’s AltarC84 and its derivatives Iride (ITA), DBA-Aurora (AUS) and Saragolla (ITA) (**Supp. Info Table 1**). Interestingly, the connection between Hap-A1 and Hap-Hap-A4/Hap-A6/Hap-A7 is represented by Hap-A5 and Hap-A2. In particular, Hap-A5 is composed by three WEW and one DEW, while Hap-A2 by five DEW and one WEW. These two haplotypes are similar, with SNP differences at 1,251 bp (G/T) and 1624 bp (A/T) after the beginning of ONT sequencing. These two haplotypes could have played a role in the origin and differentiation of *GNI-A2* related haplotypes, such as Hap-A4/Hap-A7 (five and four DEW respectively) and Hap-A6 (four DWC). Hap-A6 differs from the other two haplotypes for a 3019 bp deletion from 1,705 bp to 5,453 bp, responsible for the *GNI-A2*-specific haplotype and increased GNS. Furthermore, Hap-A6 is closely related to DEW haplotypes Hap-A4 and Hap-A7, which differ by only one SNP mutation (C/G) at position 5,603 bp (**Supp. Info Table 1**).

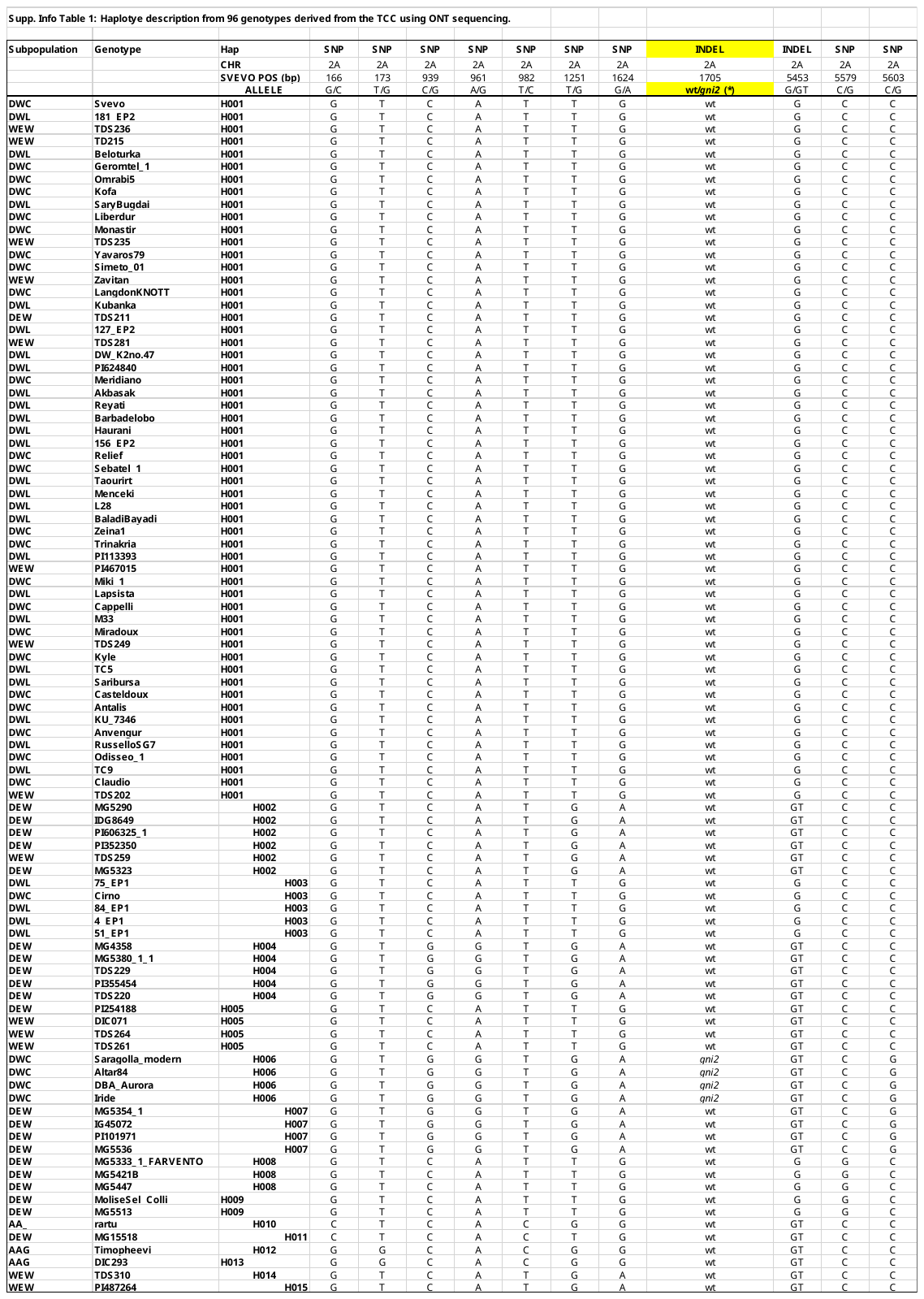

To monitor the frequency and distribution of the AltarC84 Hap-A6 *GNI-A2* deletion allele in modern durum cultivars, we then inspected its frequency in an elite worldwide durum cultivar panel (‘Durum Panel UNIBO’) of 211 durum cultivars registered between 1974 and 2000 and evaluated at CIMMYT (Obregon) for grain yield and grain yield components under optimal growth conditions (**Suppl Data S5-6**). Local SNP haplotypes and GWAS analysis confirmed that the Hap-A6 of *GNI-A2* was only identified in cultivars while it shows a significant association with GNS, particularly in the narrow interval delimited by IWB32396 and IWB32205, including the *GNI-A2* gene (**Suppl Data S4**). Interestingly, this narrow interval proved to be highly diagnostic and identity-by-descent (IBD) with AltarC84 (IWB32396-IWB32205) and in durum cultivars. SNPs outside of this IBD-region (IWB32396-IWB32205) of AltarC84 showed a lower association with the phenotype, reducing the power to detect haplotypes associated with the high-fertility floral trait (**Suppl Data S5-6**). Hap-A6 was then traced in a more recent set of 456 cultivated advanced durum lines representative of the CIMMYT breeding program. The AltarC84-derived deletion Hap-A6 was detected in 86 lines out of 456, based on a genotyping analysis using KASP markers tagging *GNI-A2* haplotype (from KUBO45 to KUBO 50) and the PCR-specific assay (STS), showing an increase to a frequency of 19%, consistent with the favorable allele effect on grain yield (**Suppl Data S7-8**). The CIMMYT breeding lines were characterized by grain yield (GY) and thousand grain weight (TGW) in different field trial managements, such as full irrigation and stress conditions (reduced irrigation-drought and heat) (**Suppl Data S8**). By only comparing genotypes with or without *gni-A2_del_* for GY and TGW, we found that GY is on average significantly increased in genotypes carrying the *gni-A2_del_* high-fertility allele in irrigated field trials. Importantly, the presence of this high-fertility allele was not detrimental under stress conditions (**Supp. Info Figure 2**). Moreover, on average this mutant allele reduced TGW significantly under all trial conditions; however, this clearly demonstrates that the *gni-A2_del_* deletion allele still has a positive effect on total grain yield, even if grain size and TGW are slightly decreased (**Supp. Info Figure 2**).

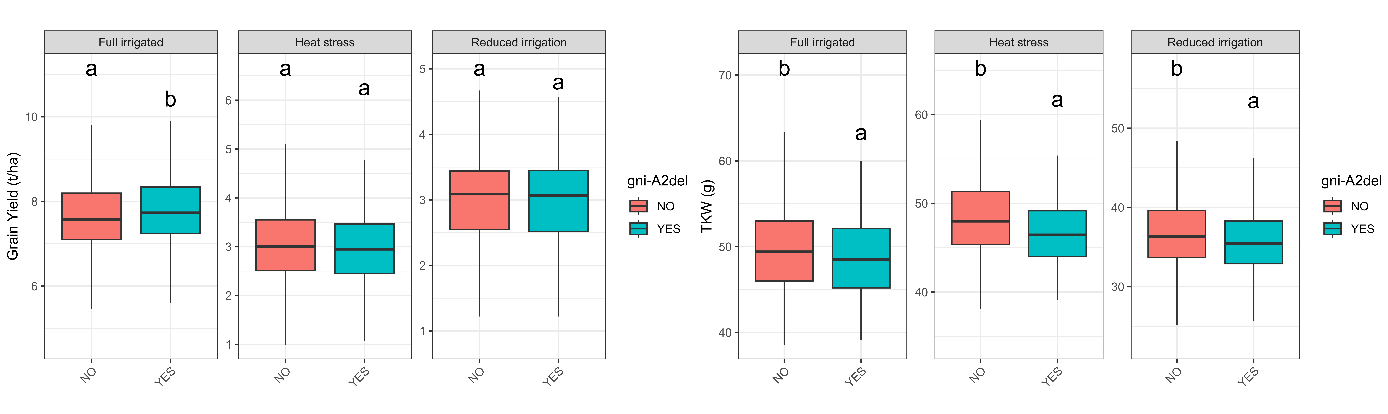
**Supp. Info Figure 2**: Variation for grain yield and TGW between durum wheat genotypes carrying *gni-A2_del_* and WT alleles in different growth conditions. Trials for grain yield and TGW were conducted under full irrigation, heat stress, and reduced irrigation-drought. A Tukey HSD test was performed using *p*-value <0.05 as a statistical significance threshold.

To shed more light on the origin of the AltarC84-derived deletion Hap-A6, we further monitored the haplotype configuration at *GNI-A2* in a wider sample of the tetraploid TGC collection^2^, including 280 DWL, 175 DEW, one spelt, and 48 DEW (**Suppl Data S9**). From this analysis, it is evident that the AltarC84 can be traced back directly to domesticated emmer (DEW) and wild emmer (WEW), where the AltarC84 long-range haplotype is the most frequent, particularly in DEW from India/Ethiopia and WEW from the Southern Levant, and not to the more recently evolved *T. turgidum* landraces (TWL). However, as already observed in the re-sequenced tetraploid core set of 96 accessions, the complete AltarC84-derived Hap-A6, including the *GNI-A2* deletion, post-dates the ancestral emmer long-range SNP-based haplotype, as none of the emmer accessions showed the *GNI-A2* deletion, clearly suggesting post-domestication selection.

**Describe CAPS marker development for *GNI-B2* (1 bp deletion) in a diverse population.**

We developed a diagnostic CAPS marker targeting the 1 bp indel of *GNI-B2* to further estimate the frequency of this loss-of-function allele, i.e. *gni-B2_298_del_*, among tetraploid wheat germplasm. All tested wild emmer accessions (**Suppl Table S2**) carried the functional *GNI-B2* allele, indicating that the *gni-B2_298_del_* allele most likely arose after domestication. Only during the selective breeding of durum cultivars, the frequency of the *gni-B2_298_del_* allele significantly increased to ~45% of the durum accessions (**Suppl Figure S2E**).

**Brief details on the field experiments and grain yield data from GABI WHEAT and BRIWECS.**

GABI WHEAT^3^: Phenotypic data were obtained from 358 winter and 14 spring wheat varieties. However, only 197 accessions were used for our re-sequencing analysis of *GNI-A1* and *GNI-A2*/-*B2*/-*D2*. Field trials were conducted across eight environments in France and Germany during 2009 and 2010 (2009/2010 Andelu/France: 48.8°N 1.8°E, height = 120.8m; 2009/2010 Seligenstadt/Germany: 50.0°N 8.9°E, height = 113.4m; 2009/2010 Wohlde/Germany: 54.4°N 9.2°E, height = 11.6m; 2010 Janville/France: 48.2°N 1.8°E, height = 135.0m; 2010 Saultain/France: 50.3°N 3.5°E, height = 72.5m). Field experiments were conducted using an alpha-lattice design with two replications.

BRIWECS^4^: Field trials were conducted in full-sized grain yield plots (harvested area 4.5–12 m^2^ depending on site-specific sowing and harvesting machinery), across a total of seven locations throughout Germany (**Suppl Table S3**), characterized by diverse soil conditions (Supplementary Table 3), in four consecutive growing seasons from 2014–2015 to 2017–2018. Plots were sown with a sowing density of 330 viable grains per meter square. The main field trials, in which all 191 cultivars of the panel were tested, were performed at each of six locations over the growth seasons 2014–2015 and 2015–2016. In each of the 12 year × location environments in the main trials, the 191 cultivars were grown in at least two replicates, sown side by side under each of the three different cropping intensities, designated as HiN/HiF, HiN/NoF and LoN/NoF treatments. The HiN/HiF treatment received mineral fertilizer at a total nitrogen supply rate of 220 kgN ha^−1^ (fertilization adjusted for soil mineral nitrogen, N_min_) along with full intensity of fungicides, insecticides and growth regulators, representing standard agrochemical applications under intensive wheat production conditions in western Europe. The HiN/NoF treatment also received a total nitrogen supply of 220 kgN ha^−1^; however, no fungicides were applied. The LoN/NoF treatment was supplied with only 110 kgN ha^−1^ and no fungicides were applied. Full details of sowing dates are shown in Supplementary Table 3 of the paper from above.
